# Cross-species proteomics quantification pipeline distinguishes donor versus host extracellular matrix in explanted biomaterials

**DOI:** 10.1101/2025.06.24.661315

**Authors:** Rachel M.E. Cahalane, Bart Meuris, Joan T. Matamalas, Mark C. Blaser, Marie Billaud, Taku Kasai, Amber Hendrickx, Ludger J.E. Goeminne, Jochen Muehlschlegel, Filip Rega, Masanori Aikawa, Laoise McNamara, Sasha A. Singh, Cassandra L. Clift, Elena Aikawa

## Abstract

Xenogenic biomaterial durability, including bioprosthetic heart valves (BPVs), is compromised by pathological extracellular matrix (ECM) remodeling, resulting in progressive structural degeneration. Mass spectrometry-based proteomics can help reveal BPV degeneration mechanisms; however, peptide sequence similarity between donor and host species complicates protein-level analysis. We present a cross-species proteomic analytical strategy for xenogenic biomaterials and cross-species proteomic datasets. *In silico* tryptic digestion of human and bovine protein databases identified over 400 overlapping proteins with a high protein percent identity. Explanted human BPV tissue was divided into macroscopically distinct regions of degeneration and analyzed by mass spectrometry. A peptide-level strategy quantified protein abundances in a species-delineated analysis. We highlighted degeneration region-specific depositions of key human ECM proteins and bovine ECM proteins whose abundance is time dependent. We demonstrated that single-species analysis of a cross-species proteome results in inaccurate quantification. This study highlights the importance of distinguishing between donor and host species proteomes for accurate protein quantification. While focused on clinically explanted biomaterials, our approach is broadly applicable to all forms of xenotransplantation and the use of xenogenic matrices.

## Introduction

The naturally occurring extracellular matrix (ECM), a dynamic structural scaffold of proteins and proteoglycans, is a promising biomaterial for applications ranging from decellularized organs and tissue sections to purified proteins^1^. In cardiovascular applications, the most common use of xenogenic-based biomaterials is bioprosthetic heart valves (BPV) constructed from glutaraldehyde-fixed bovine pericardium^2^. However, hemodynamic deterioration of BPVs impacts their durability. Circulating and cell-derived proteolytic enzymes^3^, or external stimuli including cyclic mechanical loading^4^, contribute to the degradation of the collagen-rich xenogenic donor matrix post-implantation. Concurrently, BPV adsorb host circulating proteins^5^ and the host synthesizes and deposits new ECM onto the xenogenic matrix^6^. This dynamic remodeling process limits their lifespan, almost one-third of BPV exhibit deterioration at 8 years follow-up^7^, and most require revision within 15 years^8^. To extend BPV lifespan, it is crucial to understand post-implantation remodeling, which drives BPV failure and involves ECM components from both donor and recipient species. Furthermore, ongoing donor shortages continue to propel experimental research in xenotransplantation, where xenogenic-derived ECM is exposed to a human host^9^.

To better understand the ECM in various physiological and pathological contexts, mass spectrometry (MS)-based proteomics has become the preferred method for profiling the ECM^10^. The matrisome has been defined as the ensemble of all ECM-related proteins and includes ECM core components such as collagens, glycoproteins, proteoglycans, and matrisome-associated (ECM-affiliated proteins and regulators)^11^. MS has enabled the identification and quantification of over 1,000 proteins in both the human^12^ and bovine^13^ matrisomes. MS-based proteomics has been recently applied to explanted degenerated BPV leaflets^3,14^, but the cross-species aspect has yet to be addressed.

In traditional bottom-up MS-based proteomics, proteins are digested into peptides prior to liquid chromatography-tandem MS analysis. Fragmented peptide spectra are then searched *in silico* against amino acid sequence databases for identification. In cross-species proteomics, including explanted xenogenic biomaterials which contain both donor and host proteins, reference databases for both species are required. Given the 80% genetic similarity^15,16^ between the *Homo sapiens* and *Bos taurus* genomes, the similarity of peptide sequences between species may complicate downstream protein identification and quantification. Sufficient sequence coverage over divergent residues can theoretically differentiate species proteins. In practice, however, the protein grouping and quantification of conserved peptide sequences must be given careful consideration.

Previous cross-species MS-based proteomic preclinical research has mainly relied on peptide-level quantification: human-mouse tumour xenografts^17^ and serum markers^18^, and human collagen hydrogel injection into a mouse infarcted heart^19^. Protein-level quantification methods have utilized serial species-specific search strategies to curate custom databases for non-unique peptides only and have been used to quantify shared proteins between different strains of bacteria^20^. However, in this case, the two species were not mixed in the same sample nor acquired in the same spectra. Custom database curation based on background contaminants has been performed for multi-species *in vitro* culture preparations (enzymatic or media-derived contaminants)^21,22^. To our knowledge, species-unique peptides have not been utilized to quantify species-specific protein abundances within a mixed-species proteome in xenogenic biomaterial applications. Such a cross-species protein-level analysis is especially critical for understanding explanted xenogenic biomaterials and potential mechanisms of failure. Recently, novel proteomic quantification packages have been developed that utilize combinatorial approaches for protein quantification that consider peptide spectral counts, summed spectral intensities, and peptide-based linear regression models^23^, however, these have not yet been used for cross-species quantifications as we have in this study.

In this study, we utilized MS-based proteomics to characterize the background proteome for xenogenic (bovine) matrix tissues explanted from human patients. We identified potential overlapping proteins between a custom *Bos taurus* and a complete *Homo sapiens* library and examined explanted BPV protein specificity. This approach enabled us to investigate both bovine and orthologous human proteins in degenerated bioprosthetic tissue subtypes, as well as varying implantation durations. Additionally, we explored how neglecting the donor species proteome impacts the quantification of host species protein intensities, emphasizing the importance of cross-species considerations to ensure accurate protein-level quantification.

## Results

### *In Silico* analysis reveals an overlap between *Bos taurus* and *Homo sapiens* protein databases, with a high percent protein identity and shared tryptic peptides

We aimed to optimize a proteomics quantification strategy which delineates between host and donor species in xenogenic biomaterials, in this case BPV (**Fig. 1a**). In bottom-up proteomics analysis, it is essential to properly assign spectra to peptides, and subsequently proteins. This is completed by searching the spectra against protein databases containing all potential amino acid sequences (**Fig. 1b**). However, there is an exponential loss of statistical power in spectral assignments with larger reference databases^24^. To overcome this, we created a custom bovine background database by processing and searching non-implanted bovine pericardium, limiting our *Bos taurus* reference database to a subset that is of analytical interest and biological relevance^25^ (**Fig. 1c**). Our custom bovine background database included 417 identified proteins, 400 of which were shared between the custom limited *Bos taurus* background and the complete *Homo sapiens* databases (overlapping proteins: 96% of custom bovine, 2% of human) (**Fig. 1d**). Protein sequence alignment of these 400 proteins revealed a high percent identity (conserved amino acid sequences, median: 87.88%) (**Fig. 1e, Supplementary Fig.1**). Twenty-one overlapping proteins had 100% identity, rendering them indistinguishable by peptide sequence, regardless of acquisition quality or analytical approach (**Supplementary Table 1**). Proteins with >85% percent identity were found to be enriched for ECM-related Gene Ontology processes, including Basement Membrane Organization (GO:0110011), Elastic Fiber Assembly (GO:0048251), and Collagen Fibril Organization (GO:1904026) (**Supplementary Fig. 2**). To consider shared peptide sequences relevant for proteomics, *in silico* tryptic digestion of the 400 overlapping proteins demonstrated a 17% overlap in peptide precursor ions between the species (31% of human and 26% of bovine). The false discovery rate threshold for determining high-confidence peptides acceptable for protein quantification is based on the peptide’s posterior error probability, which considers peptide length^26^. Shorter peptides (<12 amino acids in length) were more likely to be shared between human and bovine species, accounting for 52.26% of spectral library similarity (**Fig. 1f**).

**Figure 1:**
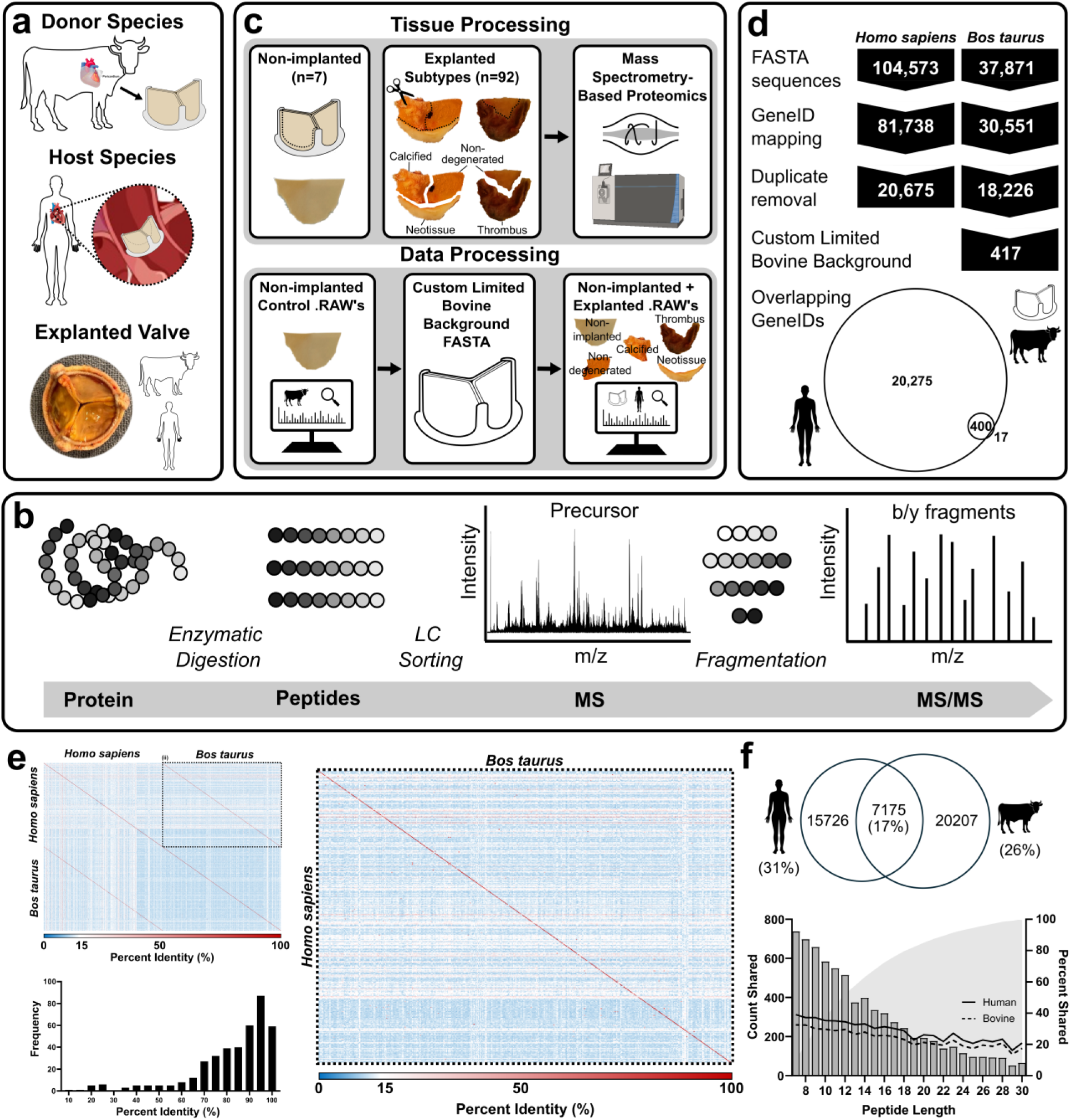
*In silico Homo sapiens* and *Bos taurus* proteomes. **a**, Explanted glutaraldehyde-fixed bovine pericardium study. Donor species (*Bos taurus*) pericardial tissue undergoes glutaraldehyde fixation and constructs bioprosthetic valve (BPV) leaflets. Bioprosthetic valves are implanted into the host species (*Homo sapiens*). Explanted degenerated BPV tissue contains donor and host species proteins. **b**, Bottom-up label-free mass spectrometry workflows consist of enzymatic digestion of proteins to peptides. Peptides are sorted by liquid chromatography (LC) followed by precursor ion detection in the MS1 spectra. Peptides are then fragmented and detected in the MS2 spectra. **c**, Non-implanted and explanted tissue processing for mass spectrometry sequencing. Gross pathological segmentation was performed on explanted BPV leaflets into non-degenerated, thrombus, neotissue, and calcified regions. Data processing involves the creation of a custom, limited bovine background file by searching the non-implanted control tissue against a complete UniProt *Bos taurus* library. The non-implanted and explanted tissues are then searched against the custom limited bovine background file and a complete Uniprot *Homo sapiens* library. **d**, *In silico* complete *Homo sapiens* and custom limited *Bos taurus* background FASTA protein overlap. **e**, Percent identity matrix (amino acid sequence comparison) for n=400 potentially shared proteins. Enlarged *Homo sapiens* versus *Bos taurus* matrix section. (Percent identity histogram for the direct species comparison cases (n=400 potentially shared proteins, diagonal red line). **f**, Venn diagram overlap of the peptide sequence acquired from *in silico* protein tryptic digestion of n=400 potentially shared proteins. Peptide duplicate count and cumulative % count (shaded area), and percent shared peptides for human and bovine species per peptide length.

### ECM-related proteins dominate orthologs

After assessing the overlap between both whole and tryptic proteomes in *Bos taurus* and *Homo sapiens* databases, we established a BPV analytical pipeline using 7 non-implanted, control and 92 explanted, representing distinct degeneration subtypes. Proteomic analysis identified 1,815 proteins in the combined non-implanted and explanted tissue search. Based on the identified peptide sequences and their degree of overlap between species, proteins were categorized by species specificity: “bovine-specific” (n=4) or “human-specific” (n=1,701) where the distinguishing peptide(s) were only identified in one species; “orthologous” (n=24), where each species’ peptides have a distinguishing amino acid feature identified; and “cannot differentiate” (n=86) where there is no distinguishing feature in the tryptic peptides identified (**Fig. 2a-b**).

**Figure 2:**
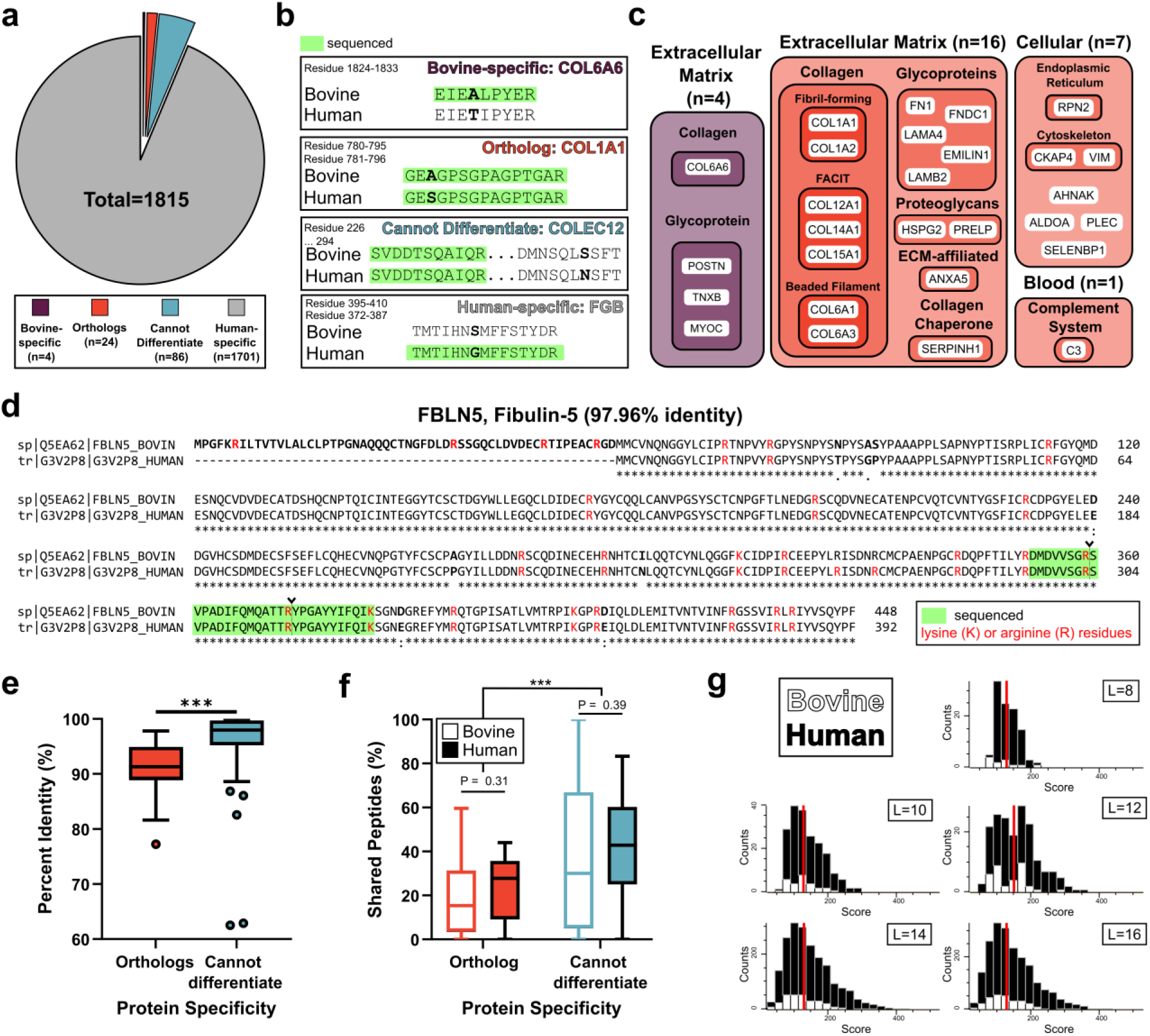
Protein and peptide specificity. **a**, Pie chart of protein specificity from the non-implanted and explanted tissue search. In total 1815 proteins were identified: bovine-specific (n=4), ortholog (n=24), cannot differentiate (n=86), and human-specific (n=1701). **b**, Representative sequenced amino acids for each specificity case (bovine-specific: Collagen Type VI Alpha 6 Chain (COL6A6), ortholog: Collagen Type I Alpha 1 Chain (COL1A1), cannot differentiate: Collectin Subfamily Member 12 (COLEC12), and human-specific: Fibrinogen Beta Chain (FGB)). **c**, Protein biological function grouping for bovine-specific and ortholog proteins. Matrisome classification was guided by reference bovine^13^ and human^12^ matrisome databases. **d**, Representative UniProt alignment for Human and Bovine Fibulin 5 (FBLN5) (97.96%). Red Lysine and Arginine residues indicate peptides. Sequenced peptides are highlighted in green. Inverted caret indicates cleavage point. **\*** Fully conserved residues. : Conservation between groups of strongly similar properties (Gonnet PAM 250 score > 0.5). . Conservation between groups of weakly similar properties (Gonnet PAM 250 score ≤ 0.5). No symbol, non-conserved residues^48^. **e**, Tukey boxplot of percent identity for ortholog versus cannot differentiate proteins. **f**, Tukey boxplot of percent shared peptides for ortholog and cannot differentiate proteins per bovine and human species. **g**, Histograms of MaxQuant peptide score for identified bovine and human peptides, by length. Red lines represent the centroids.

The bovine-specific proteins were exclusively associated with the ECM (COL6A6, POSTN, TNXB, and MYOC). Two-thirds (16/24) of the orthologous proteins were related to the ECM (collagens, glycoproteins, proteoglycans, ECM-affiliated, and ECM regulators). The remaining orthologs were related to cellular processes (7/24) or the complement system (1/24) (**Fig. 2c**). Proteins that could not be differentiated between human and bovine based on peptide sequence were related to elastic fiber assembly, cell-cell adhesion, mitophagy, and lipid localization processes (**Supplementary Table 2**). Human-specific proteins were related to immunity, cellular metabolism, basement membrane organization, and blood coagulation/fibrinolysis processes (**Supplementary Table 3**). The percent identity (**Fig. 2d**) of cannot differentiate was significantly higher than orthologous proteins (96.53% vs 91.33%, P-value < 0.0002, **Fig. 2e**). Additionally, more cannot differentiate peptides were shared compared with ortholog peptides (37.76% vs 24.39%, P < 0.0004, **Fig. 2f**), demonstrating that even with complete sequence coverage, specific proteins are unable to be differentiated between species. Peptide confidence scores also showed similar distributions between species for peptides of the same length (**Fig. 2g**).

### Bioprosthetic degeneration is heterogeneous

Explanted degenerated BPVs were segmented into degeneration subtypes to address pathomorphological tissue heterogeneity: non-degenerated (N=36), thrombus (N=17), neotissue (N=17), and calcified (N=22) (**Fig. 3**). Non-implanted control tissue (glutaraldehyde-fixed bovine pericardium) appeared macroscopically intact, with a dense collagen matrix and dispersed fibroblast-like cells. Explanted, macroscopically non-degenerated regions, no longer contained dispersed cells but exhibited some sparse cells or monolayers at the leaflet blood contacting surface. Thrombotic regions presented as a thick, dark and structurally distinct matrix overlaying or within the pericardial tissue. Surface thrombus typically contained cells with a rounded morphology. Neotissue formed exclusively on the BPV leaflet surface, as a structurally aligned and intact collagenous matrix containing spindle-shaped cells. Histological analysis also confirmed visible calcified deposits.

**Figure 3:**
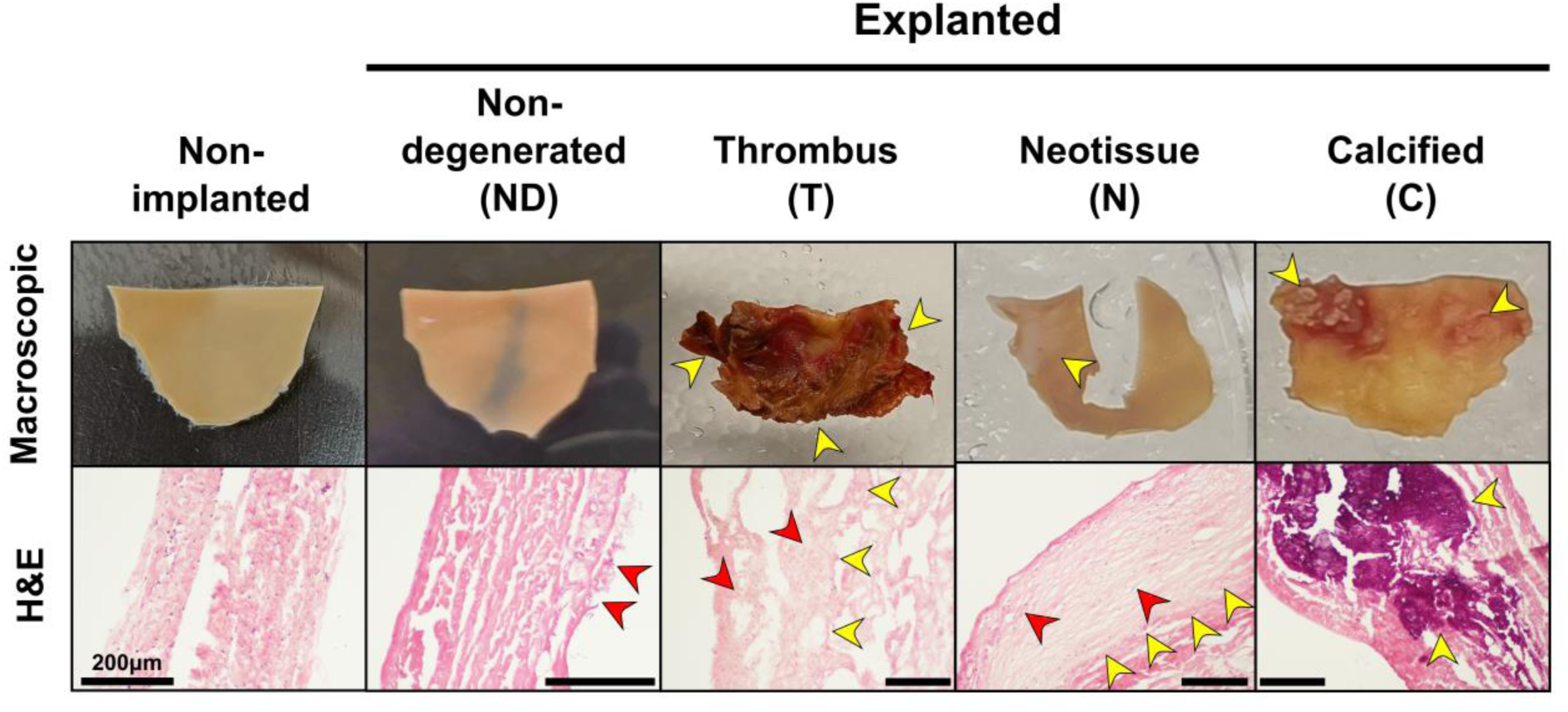
Gross pathology and histopathology of bioprosthetic (BPV) tissue. Macroscopic appearance and corresponding H&E-stained cross sections of non-implanted control bovine pericardial BPV tissue versus explanted BPV tissue subtypes: non-degenerated, thrombus, neotissue, and calcified. Yellow arrows indicate the tissue subtypes. Red arrows indicate observed cells. The H&E scale bar represents 200μm.

### Spatiotemporal profiling of cross-species proteomes reveals transition of ECM in bioprosthetic valves

We next utilized a protein quantification strategy that considers peptide spectral counts, summed spectral intensities, and peptide-based linear regression models to accurately delineate species-specific orthologous protein intensities^27,28^. This peptide-centric quantification also allows for visualization and interpretation of protein-domain-specific contributions to quantification. We did this in the context of BPV degeneration subtypes to quantify the contribution of bovine tissue degradation and human neotissue deposition. Several bovine proteins were present only in non-implanted samples, suggesting rapid degeneration in explanted tissue. Structural proteins, including COL1A1 and COL1A2, were less abundant but were present in explanted tissue. Within both species, tissue segments unbiasedly clustered into non-explanted vs. explanted, and neotissue vs. degenerated subtypes (non-degenerated vs. thrombus and calcified). However, ECM orthologs were highest for non-implanted (bovine) and explanted neotissue (human), demonstrating degradation of bovine ECM followed by deposition of new human ECM in neotissue segments (**Fig. 4a**). Human COL1A1 pre-processed peptide intensity was significantly higher than bovine in neotissue subtypes, reflecting observed matrix deposition. Bovine Prolargin (PRELP; a collagen- and proteoglycan-binding protein) intensity decreased in explants, while human PRELP levels surpassed bovine in all explanted subtypes. Bovine cellular remnants (Vimentin (VIM); a well-described marker of fibroblasts) appear in all tissues while human VIM was present in explants, consistent with cellular monolayer observations, but is highest in thrombus and neotissue subtypes (**Fig. 4b, Supplementary Fig.3**).

**Figure 4:**
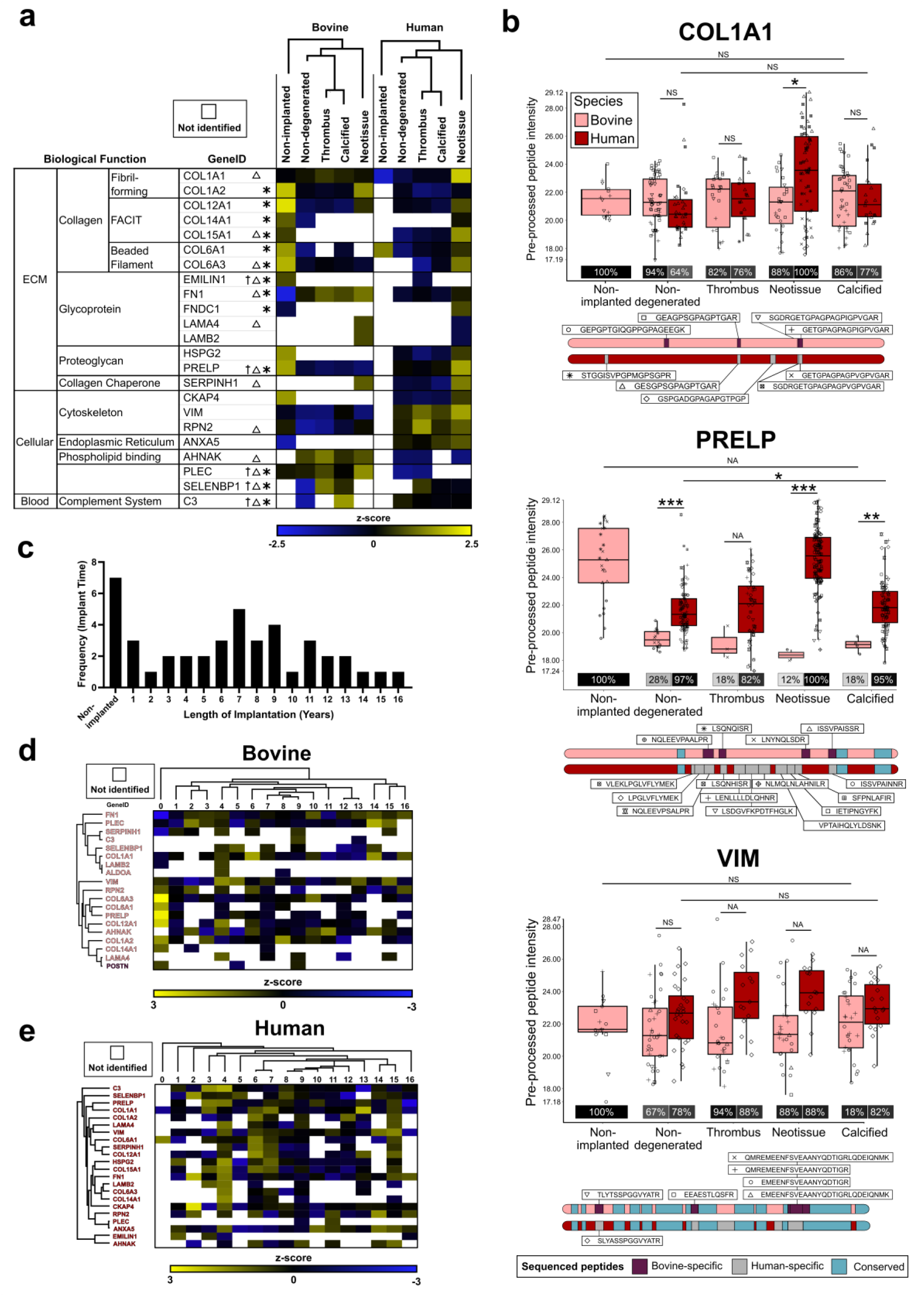
Spatiotemporal protein profiling. **a**, Heatmap of bovine proteins (specific and orthologs) according to median peptide intensity per tissue subtype. Heatmap is split by species (bovine and human). Two-way ANOVA significance: *species, Δsegment, †interaction (P-value threshold = 0.05). **b**, Bovine and human ortholog pre-processed MSqRob peptide intensities for Collagen 1A1 (COL1A1) and Prolargin (PRELP). Percentage greyscale boxes indicate the percentage of study samples that presented with these peptides. Sequenced peptide site mapping onto protein structure for bovine and human counterparts. Blue, purple and grey regions indicate conserved, bovine-specific, and human-specific peptides, respectively. Imputed human COL1A1 peptides for non-implanted samples were excluded. Statistical significance: *** P ≤ 0.0005, ** P ≤ 0.005, * P ≤ 0.05, NS = non-significant, NA = not applicable (no matching peptides). **c**, Overall histogram for the number of donors per length of implantation. Note: length of implantation details for one donor is unavailable. **d-e**, Heatmap of bovine (d) and human (e) proteins per implant duration (years, order constrained). Proteins are unbiasedly clustered.

Samples in this study were implanted up to a median of 7.50 years before surgical removal (**Fig. 4c**). When we examined orthologs according to time-dependent behavior, bovine structural collagens (COL1A1, COL6A3, and COL1A2) and cellular remnants (VIM) were observed across all time lengths (**Fig. 4d, Supplementary Fig. 4**) Of note, bovine COL1A1 intensity peaked after 6 years of implantation. Broadly, bovine proteins clustered into 1-3, 4-13, and 14-16 implantation duration (years, order constrained). Human samples showed less grouped clustering, but many human proteins peaked in intensity between 2 – 7 years of implantation.

### Network prioritization of proteins with bovine degradation and human deposition

While orthologous protein profiles can be visualized across degeneration subtypes and implantation duration, this approach may overlook proteins with species-specific abundance patterns. We used similarity-based metrics to prioritize proteins proteins showing distinct profiles across degeneration subtypes or species. Here, we created vector matrices, where each vector represents a protein, and each vector dimension represents a species-tag (**Fig. 5a**). By calculating the distance between these vectors and using that distance as an edge weight in a protein-based network, we can identify which proteins have the most divergent species-tag patterns, indicating the greatest bovine degradation and subsequent human deposition. When analyzed independent of degeneration subtype, COL1A1, COL6A3, AHNAK, and VIM showed the highest species-based protein dissimilarity within any tissue subtype (**Fig. 5b**). These proteins, in addition to COL12A1, also have a large dissimilarity across degeneration segments, independent of species specificity (**Supplementary Fig. 5**), suggesting that distinct bovine degradation and human deposition profiles significantly contribute to bioprosthetic pathological degeneration.

**Figure 5:**
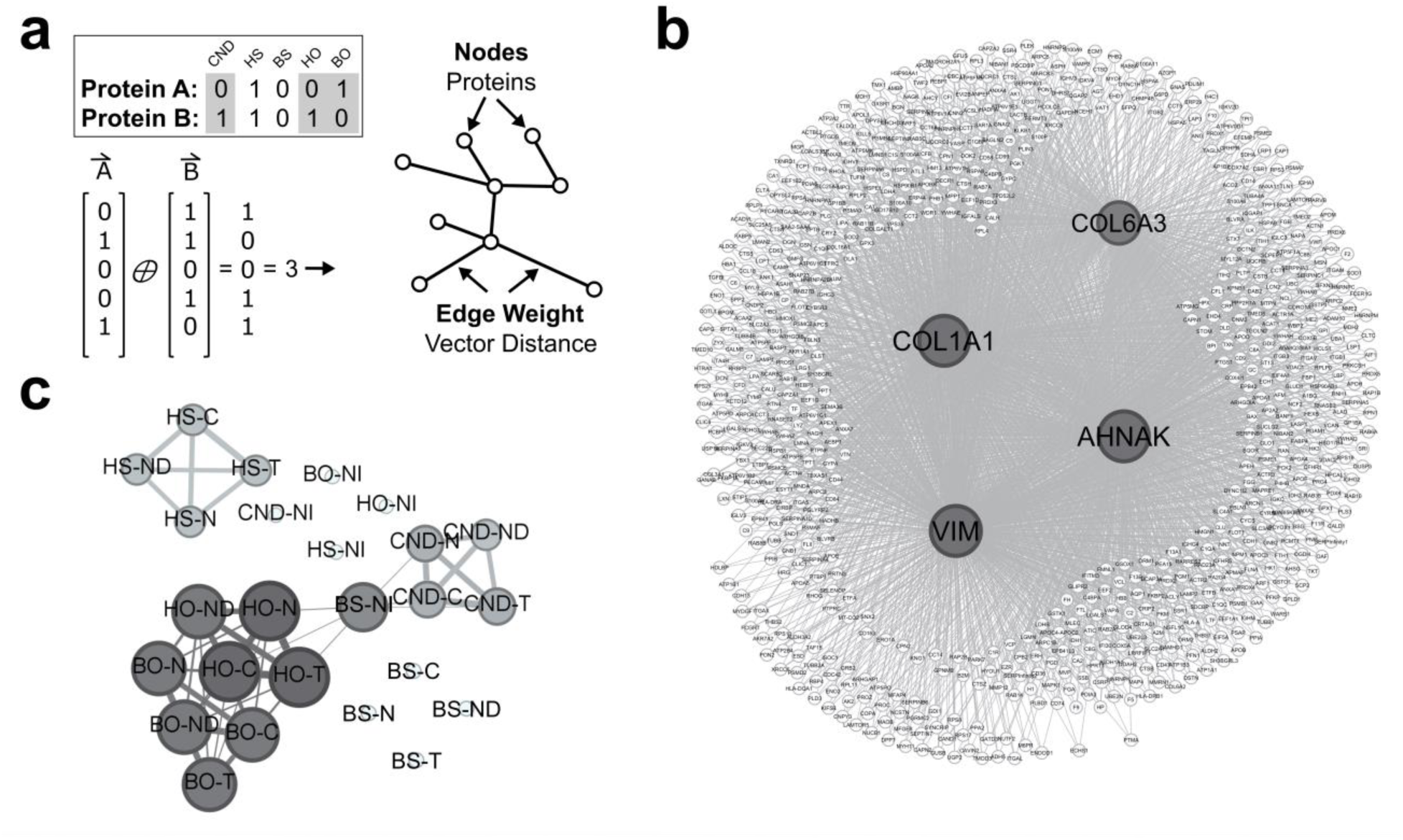
Similarity-based metrics to prioritize proteins. **a**, Protein vector matrices were constructed with 25 distinct dimensions, comprising five potential species tags (bovine-specific, bovine ortholog, human ortholog, human-specific, and “cannot differentiate”) across five explanted BPV tissue subtypes (non-implanted, non-degenerated, thrombus, neotissue, and calcified). Zero or 1 represents unidentified and identified per species-degeneration combination. The distance between each protein vector in the protein-protein interaction network was calculated and used to weigh the network. **b**, Hamming distance vector analysis was used to assess dissimilarity across species tags (bovine and human), which were then used as edge weights in the protein similarity network. **c**, The Jaccard Similarity Index was used to measure the overlap in species-tag presence between proteins across the 25 distinct dimensions. (HS = human-specific, BS = bovine-specific, BO = bovine ortholog, HO = human ortholog, CND = cannot differentiate) and tissue subtype (NI = non-implanted, ND = non-degenerated, T = thrombus, N = neotissue, C = calcified). Dissimilarity and Similarity thresholds were set below or above 0.5 vector distance, respectively.

We also used this vector distance-based network analysis to identify the greatest overall protein similarity between species-degeneration combinations across all 25 dimensions (protein specificity and degeneration subtype). Neighboring species-degeneration nodes showed the greatest similarity in total proteome abundance. Human-specific proteins showed distinct profiles from bovine-specific, orthologs, and cannot differentiate proteins, but were unaffected by tissue subtype, implying a baseline infiltration level with abundance (rather than presence) changing during bioprosthetic degeneration. Bovine-specific proteins in non-implanted BPV proteome bridge cannot differentiate and orthologs. Bovine-specific protein detection is dissimilar across degeneration segments, indicated by a lack of edge-connectivity, suggesting stage-specific bioprosthetic matrix degradation (**Fig. 5c**).

### Single-species search of a cross-species proteome leads to data misinterpretation in a degeneration subtype-specific manner

To assess the impact of the conventional single-species search on cross-species samples, we analyzed our explanted BPV tissue spectral files against a human FASTA alone and compared the results against our cross-species analysis (**Fig.6a; Supplementary Fig. 6)**. Confidently identified proteins were largely preserved between the two searches (94%) (**Fig. 6b**). Proteins previously classified as orthologs and cannot differentiate in the cross-species search were still identified in the human-only search (96% and 95%, respectively; **Fig.6c**). No bovine-specific proteins (COL6A6, POSTN, TNXB, and MYOC) were identified in a human-only database search. The quantification impact was assessed by comparing the median abundance rank of proteins in the human-only search to the cross-species search (**Fig. 6d, Supplementary Fig. 7a**). In neotissue and thrombus, the under- and overestimated protein groups account for as much as 26.45% and 27.22% of the measured proteome, respectively. In the case of neotissue segments, orthologous collagens COL1A2, COL1A1, and COL6A3 were underestimated in the human only search (rank 1189, 912, and 1408) compared with the cross-species search (rank 289, 204, and 320). The under- and over-estimated proteins are also specific to each tissue subtype, with ≤2% shared (**Fig. 6e**). We show specific biological processes enriched in over- and under-representative data including leukocyte differentiation, lipid particle remodeling, platelet aggregation, and coagulation, which are critical pathways implicated in valve pathogenesis in the context of thrombosis and calcification (**Supplementary Fig. 7b**). Species-delineated neotissue COL1A1 comparisons underscore the importance of cross-species considerations and the risks of ignoring multi-species effects in proteomics (**Fig. 6f**).

**Figure 6:**
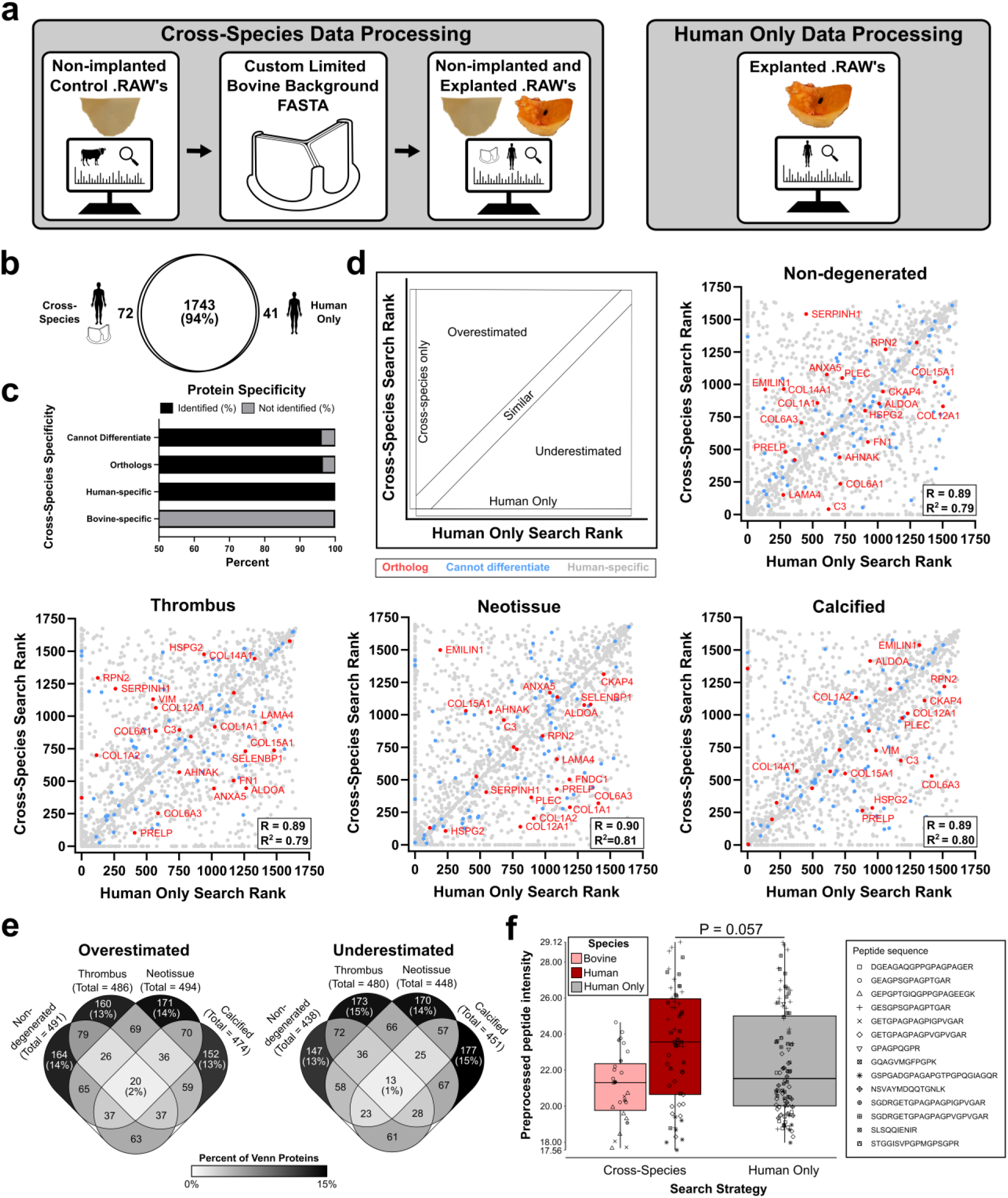
Comparison of cross-species search results against a human-only search. **a**, Human-Bovine cross-species data processing consists of the creation of a custom limited bovine background file by searching the non-implanted control tissue against a complete UniProt *Bos taurus* library. The non-implanted and explanted tissues are then searched against the custom limited bovine background file and a complete UniProt *Homo sapiens* library. Human-only data processing includes searching the explanted tissues only against a complete UniProt *Homo sapiens* library. **b-c**, Comparison of protein identifications and specificity between the two searches. **d**, Median abundance rank comparisons between the cross-species and human-only searches for non-degenerated, thrombus, neotissue, and calcified tissue subtypes. Scatterplot region analysis: X = 0 (Cross-Species Only), Y = 0 (Human Only), X = Y ± 100 (Similar), x > y (Overestimated), y > x (Underestimated). Rank of 1 = highest median abundance. Note: Bovine-specific proteins were removed from the cross-species rank comparisons. **e**, Venn diagrams of the overlapping over- and underestimated proteins per tissue subtype. **f**, Cross-species versus human-only search pre-processed MSqRob peptide intensities for Neotissue Collagen 1A1.

## Discussion

Xenogenic biomaterials, such as BPVs, have limited durability due to ECM pathological remodeling. While MS-based proteomics is a valuable tool for identifying molecular mechanisms of ECM degeneration, protein quantification is complicated by peptide sequence similarity between species in multispecies datasets. Here, we presented a proteomic data processing pipeline specific for analyzing two species (bovine and human), in the context of BPV failure. We identified key ECM constituents with bovine-specific degradation and human-specific deposition in a spatiotemporal manner. We also demonstrated how disregard for the cross-species component in the search strategy changes the relative abundance-based rank of proteins in a degeneration subtype-specific manner, under- and over-estimating changes in key biological processes linked to thrombosis and calcification.

The collagen-rich matrix of glutaraldehyde-fixed bovine pericardium preserves mechanical integrity, making it suitable for cardiac devices under cyclic loading^2^. Given their known abundance in pericardium^29^ and involvement in both fibrosis^30^ and the foreign body response^31^, we hypothesized that ECM-related proteins would be prominently represented among the orthologs identified in the current study. Here, COL1A2, COL6A1, COL6A3, and COL12A1 and ranked in the top 5 non-implanted BPV proteins sorted by median abundance. Previous glutaraldehyde-fixed bovine pericardium immunoproteomic studies have identified xenoantigens^32,33^. Up to 133 different proteins were identified that either elicited a humoral immune response with rabbit sera^33^ or were purified using affinity chromatography^32^, underscoring the potential host response to the xenogenic ECM^45^. Proteins exhibited diverse functions, with 74% localized to secretions, membrane, cytoplasm, or matrix. Similarly, 67% of the proteins in the current study were localized to the same subcellular regions.

Before the publication of the bovine matrisome^13^, previous proteomic studies of explanted BPVs were focused on human contribution to degeneration, and did not explore cross-species effects^3,14^. Abramov and colleagues profiled 24 glutaraldehyde-fixed bovine pericardium explants from patients with or without metabolic syndrome.^14^ The authors noted the contribution of ECM proteins to degeneration, particularly for metabolic syndrome samples. Our *post hoc* analysis revealed that the 50 most abundant differentially expressed proteins in explants from patients with and without metabolic syndrome had a median protein percent similarity of 91.88% with their bovine counterparts. From this list, FLOT1, RPL27, RPL8, and UBE2L3 have 100% identity across human-bovine species, preventing donor–host distinction. In comparison with our study, 7/50 proteins were present in our custom limited background file (CTSD, FABP5, FLOT1, LAMC1, LAMP1, LTBP2, and TAGLN). In a separate study, Kostyunin and colleagues used proteomics to evaluate bovine- and porcine-derived BPVs explanted from humans, and similarly used human-only databases in their search methods^3^. Three out of the 75 unique or increased proteins in BPVs compared with native valves identified were also present in our custom limited background file (STOM, APOE, and APOC3). Neither of these studies^3,14^ used non-implanted control bioprosthetic tissue in their proteomic pipelines. We have demonstrated that careful consideration of donor species proteomes is required for accurate protein quantification across bioprosthetic degeneration subtypes.

One rationale for searching a cross-species proteomic dataset against a human-only reference library is potential inconsistencies in library completeness across species. However, recent advancements in multi-omics sequencing and bioinformatics show high completeness across commonly used species. The UniProt Complete Proteome Detector score shows *Bos taurus* (bovine) has high completeness, with only 1.9% of the proteome defined as missing^34,35^, compared to other species in the same taxonomic class. Further, inclusion of all species present in the sample reduces the number of decoy PSM matches, lowering overall FDR threshold, and resulting in more confident identifications for both species^25^. Considering the ECM, the complete matrisome compendium contains just over 1000 proteins, for both *Bos taurus*^13^ and *Homo sapiens*^12^. Given the recent advancements to database completeness^10^, we and others have shown that considering only one species in database searching is a greater detriment to data quality than including both species with minimally differing database completeness^25^. This is particularly relevant for explanted xenogenic biomaterials, as we have demonstrated here for BPVs.

We identified several bovine-specific peptides contributing to relatively high corresponding protein abundance in the non-implanted bovine pericardium, which were not identified in all of the explanted segments. These proteins were largely associated with non-structural ECM, including collagen alpha-1(XV) (COL15A1), elastin microfibril interface-located protein 1 (EMILIN1), fibronectin type III domain containing 1 (FNDC1), and perlecan (HSPG2) or intracellularly localized proteins including annexin A5 (ANXA5) and cytoskeleton-associated protein 4 (CKAP4). We did not identify many glycoproteins: however, glutaraldehyde fixation does not preserve glycoproteins or native pericardial cells^36^, consistent with our histological observations.

Glutaraldehyde fixation renders bovine pericardium resistant to rapid enzymatic degradation^29,37^; however, molecular-level collagen damage can occur after cyclic mechanical loading,^38^ or through plasma-derived MMP9 proteolytic degradation^3^. Consistent with this, our data indicates higher z-score abundances of bovine COL1A1 in explanted samples compared to non-implanted controls (6 years, 14 years). Additionally, temporal analysis of human orthologs’ z-score abundances suggests a bimodal distribution. The first cluster of higher abundances occurs between 2- and 7-years post-implantation. Samples requiring earlier revision may exhibit elevated levels of human tissue remodeling, potentially contributing to the need for expedited reintervention.

The tryptic proteomic workflow used here, while capable of identifying ECM proteins, is not optimized for the robust profiling of all constituents of the matrisome. Sequential extraction of proteins based on solubility^12^ can be used, but there are inherent challenges with targeted ECM proteomics, such as high-degree of intramolecular crosslinking, post-translational modifications, and reduced accessibility of tryptic digestion, amongst other factors^12,39,40^. Moreover, the bovine xenogenic matrix was crosslinked with glutaraldehyde. We tested additional enzymatic digestion and antigen retrieval steps, but they did not increase our acquired bovine proteome (**Supplementary Fig. 8**). Therefore, no additional steps were taken in the final workflow to release crosslinked collagen for sequencing.

Our analysis of spectral libraries from *in silico* digestion showed that peptides shorter than 12 amino acids contribute to over half of the peptide sequence overlap – that is, the longer the peptide, the more likely it is to be unique. However, these peptides represented less than a third of all sequencing events in the acquired bioprosthetic tissue proteomic data. This is based on sequence overlap and does not consider charge state and fragment ion matching across peptides to identify unique features. Although not optimized in this study, this analysis suggests that excluding shorter peptides from quantification in cross-species analysis could improve data deconvolution between species. Alternatively, if there is a primary interest in one species, the search can be performed against all relevant proteome databases and then recalculate the False Discovery rate based on the species of interest.

Finally, while we demonstrated this cross-species label-free proteomic quantification method being used for BPV explants, many other potential applications exist. Xenogenic scaffolds are used in a variety of clinical applications, such as the use of porcine small intestinal submucosal-derived materials for repair of intracardiac, aortic, pericardial, and great vessel defects^41^, urinary conduit, gastric mucosal repair, as well as tendon and bone regeneration^42^. Outside bioprosthetics, patient-derived xenografts are a cross-species model widely used in cancer research to investigate tumour heterogeneity and *in vivo* therapeutic screening^43^. Additionally, fecal proteomics, useful for biomarker discovery and immune profiling of gastrointestinal tract diseases, has inherent complications due to protein contributions from the host, microbiome, and ingested food^44^. Overall, this novel analytical pipeline improves upon label-free quantification techniques currently used in cross-species proteomic analyses and has a wide array of potential applications in biomedical engineering and biomedical sciences.

In summary, this study highlights the critical importance of distinguishing donor and host species proteomes to achieve accurate protein-level quantification. Although our focus is on bioprosthetic heart valve replacement, our methodology is universally applicable to all cases of xenotransplantation or the use of xenogenic matrices.

## Online Methods

### Tissue Acquisition

Control, non-implanted bioprosthetic tissue was acquired from products non-eligible for clinical use (beyond expiration date commercial valves, Edwards Lifesciences) or donated commercial-grade material for research purposes (Boston Scientific). Control non-implanted bioprosthetic material was kept in 1) low-concentration glutaraldehyde solution (0.6%) at room temperature or 2) stored dry at -80°C. Explanted bioprosthetic valve leaflets were obtained from revision aortic valve replacement surgeries following institutional review board protocols Body/Muehlschlegel (2021P001077) and Meuris (S67924). Surgical donor inclusion criteria included hematocrit >25%, patient age between 20-85 years, aortic position, and glutaraldehyde-fixed bovine pericardial tissue. Patients with endocarditis were excluded. Collection of explanted tissues included written informed consent from all donors. All explanted tissues were either 1) placed in high-glucose Dulbecco’s modified Eagle’s medium on ice, then macroscopically segmented (see below) and stored at -80°C or 2) underwent immediate storage at -80°C.

### Pathological Tissue Segmentation

Gross pathological segmentation was performed on explanted BPV leaflets to address visual variations in degeneration according to the following macroscopic morphological guidelines: non-degenerated, thrombotic, neotissue, and calcified. These tissue degeneration subtypes were selected based on previous histological reports of explanted bioprosthetic leaflets^46,47^ in combination with our macroscopic observations of non-degenerated, thrombus, neotissue, and calcified. Non-degenerated regions were largely preserved (macroscopically) with no evidence of thrombus, neotissue, or calcification. Thrombotic regions appeared thicker and and ranged in color from bright to dark red-brown. Neotissue regions presented a distinct new surface layer of tissue with a color different to the underlying BPV leaflet, including areas of pannus overgrowth. Calcified regions contained visible mineralized deposits that were rigid upon probing.

### Histology

During the macroscopic segmentation process, representative longitudinal (tip to base) cross-sections were cut out, washed, and embedded in OCT compound (Tissue-Tek). Either one cross-section was acquired per leaflet if it included all segments defined for that leaflet/donor or per segment to perform validation on the macroscopic segmentation process. OCT blocks were stored at -20°C. Cryosections (7 μm) were prepared for histological staining using a cryostat (Leica CM3050 S). The overall morphology of the tissue was examined using Hematoxylin & Eosin (H&E) staining. In brief, H&E staining was performed by drying sections for 30 mins, followed by post-fixation in 10% formalin for 10 mins. After rinsing, sections were stained in Harris hematoxylin for 2 mins, dipped in 1% acetic acid, incubated for 1 min in ammonium water, rinsed in 70% alcohol, dipped in 1% alcoholic eosin with agitation, and dehydrated in 95% and 100% ethanol, and xylene before coverslipping in SHUR/mount (VWR). All macroscopic segmentation subtypes were confirmed by a cardiovascular pathologist in a blinded manner.

### Tissue Processing

Segmented tissue samples were kept frozen and processed into ∼ 1mm^3^ pieces using a blade. Tissue pieces were pulverized in liquid nitrogen using a pestle and mortar and were then resuspended in RIPA buffer (ThermoFisher) supplemented with protease and phosphatase inhibitors (Millipore Sigma) to extract the protein. Samples were incubated on wet ice for at least 1 hr with intermittent vortexing and were then subject to 10 mins of sonication in a benchtop ultrasonic water bath (Branson 5510). Samples were centrifuged at 2000 g for 5 min. The supernatant was collected and transferred to a new tube. The protein concentration was determined using the Bicinchoninic acid (BCA) assay on a Nanodrop spectrophotometer reading at 562 nm against a standard curve. Samples were stored at -80°C for liquid chromatography-tandem mass spectrometry (LC-MS/MS) (see below).

### Mass Spectrometry

All samples were prepared for mass spectrometry-based proteomics using PreOmics x96 iST kit according to the manufacturer’s guidelines. For non-implanted control bovine tissue and explanted BPV tissue 4 μg and 10 μg were processed, respectively. Proteins were lysed, reduced, and alkylated by heating in lyse buffer at 95 °C for 10 min. Samples were digested for 2 h at 37 °C. Peptides were purified through a cartridge, and the purified peptides were eluted off using kit solutions. Peptides were dried in a speed vacuum at 45 °C for 45 min or until dry. Samples were stored at -80 °C until the time of mass spectrometry sequencing. Some modifications for low abundance samples were performed. Deviations from standard protocol are as follows: 1) 10 μL of LYSE buffer and 2) 10 μL of Lys-C/trypsin DIGEST solution were added to all samples. Upon defrosting, 42μl of LC-LOAD solution (PreOmics iST kit) was added to the dried peptides. Samples were centrifuged at 16,000g for 1 min, and then 40 μL was aspirated and separated into a new tube. Peptide yield was quantified by Qubit assay on a Qubit 2.0 Fluorometer (Invitrogen). Samples were diluted to 31.25ng/μl using the PreOmics kit solution (LC-LOAD).

Mass spectra were acquired on Orbitrap Fusion Lumos coupled to an Easy-nLC1000 HPLC pump (Thermo Fisher Scientific). Injection volumes of 6 µl (total 187.5 ng peptides) were separated using a dual column set-up: an Acclaim™ PepMap™ 100 C18 HPLC Columns, 75 µm X 70 mm (Thermo Fisher Scientific, Cat# 164946); and an EASY-Spray™ HPLC Column, 75 µm X 250 mm (Thermo Fisher Scientific, Cat# ES902). The column was heated at a constant temperature of 45°C. The gradient flow rate was 300 nl/min from 5 to 21% solvent B (0.1% formic acid in acetonitrile) for 50 minutes, 21 to 30% solvent B for 10 minutes, and another 10 minutes of a 95%-5% jigsaw wash. Solvent A was 0.1% formic acid in mass spectrometry-grade water. The mass spectrometer was set to 120,000 resolution, and the top N precursor ions in a 3 second cycle time (within a scan range of m/z 375-1500; isolation window, 1.6 m/z) were subjected to collision-induced dissociation (CID, collision energy 30%) for peptide sequencing. The sample injection order was randomized.

### Custom limited background bovine file creation

The acquired peptide spectra for the non-implanted control glutaraldehyde-fixed bovine pericardium material (n=7) were searched with Proteome Discoverer package (PD, Version 2.5) using the SEQUEST-HT search algorithm against the *Bos taurus* UniProt database (Swiss-Prot and TrEMBL proteins, and Swiss-Prot protein isoforms: 37,871 sequences, updated May 2024). The digestion enzyme was set to trypsin and up to two missed cleavages were allowed. The precursor tolerance was set to 10 ppm and the fragment tolerance window to 0.6 Da. Methionine oxidation and n-terminal acetylation were set as variable modifications and cysteine carbamidomethylation as fixed modification. The PD Percolator algorithm calculated the peptide false discovery rate (FDR) and peptides were filtered based on an FDR threshold of 1.0%. The Feature Mapper was enabled in PD to quantify peptide precursors detected in MS1 but may not have been sequenced in all samples. Chromatographic alignment was performed with a maximum retention time shift of 10 minutes, mass tolerance of 10 ppm, and signal-to-noise minimum of 5. To be inclusive of all potential bovine proteins with our custom limited background file creation, peptides only assigned to one given protein group and not detected in any other protein group were considered unique and used for custom FASTA generation, as were proteins with only one peptide identified. From this analysis, a custom limited glutaraldehyde-fixed bovine pericardium FASTA list was created (453 sequences) and exported for use in the entire cohort search. The non-implanted control glutaraldehyde-fixed bovine pericardium proteome (filtered for proteins ≥ 2 unique peptides) is presented in **Supplementary Figure 9**.

### In silico overlap

The *in silico* overlap between the custom limited glutaraldehyde-fixed bovine pericardium FASTA (as described above) and the *Homo sapiens* UniProt database (Swiss-Prot and TrEMBL proteins, and Swiss-Prot protein isoforms: 104,573 sequences, updated May 2024) was determined. The protein percent identity (% of identical residues) of overlapping gene identifications was calculated using Clustal Omega (v1.2.2)^48^. *In silico* digestion was performed and subsequent spectral libraries were generated from the overlapping gene IDs in DIA-NN (v1.9.1)^49^. *In silico* digestion was performed and spectral library features were analyzed for peptide sequence overlap. Spectral libraries were sorted for precursor ions of 2+ charge states. Peptide sequences with multiple charge states were removed as duplicates.

### Data searching in MaxQuant

**1. Cross-species search:** Non-implanted control bioprosthetic leaflet material samples (n=7) and explanted bioprosthetic tissue samples (n = 92 degeneration subtypes in total (n = 37 patients, n = 48 leaflets)) were searched in MaxQuant (v2.4.14.0)^26^ against both the custom limited glutaraldehyde-fixed bovine pericardium background FASTA (*Bos taurus*) and the *Homo sapiens* UniProt Proteome database (both as described above) with a randomized decoy database and a false discovery rate of 0.01. The MaxQuant contaminant database was modified to remove bovine proteins overlapping with the custom limited glutaraldehyde-fixed bovine pericardium background FASTA (**Supplementary Table 4**). The following search parameters were used: digestion mode was set to Trypsin/LysC, carbamidomethyl was selected as the fixed modification, oxidation (M) and N-terminal acetylation were selected as the variable modifications, precursor and fragment mass tolerance were set to 20 ppm and 0.5 Da, respectively. Tissue segments (non-implanted, non-degenerated, thrombus, neotissue, and calcified) were set up as fractions 0-4, respectively. Match between fraction runs was allowed and set to 0.4 min. Outputs were filtered for marked potential contaminants and ≥ 2 unique peptides. Protein identifications and species specificity were determined from the MaxQuant Intensity outputs. Protein specificity was determined by the species identification in the Fasta headers. Quantification was performed using MSqRob (see below; 2.6.4) **2. Human only search:** To acquire a comparative human only search; only the explanted bioprosthetic tissue samples (n=92) were researched in MaxQuant (v2.4.14.0) against only the *Homo sapiens* UniProt Proteome database. The search was performed in the same manner as the cross-species search. Unique and razor peptides were used for quantification, and label-free quantification was turned on. MaxQuant ProteinGroup outputs were filtered for potential contaminants, randomized hits, and a minimum of 2 unique peptide sequences. Protein-level species specificity was determined and GeneID was assigned based on FASTA headers. Protein isoforms within the same specificity category were removed. Median abundances were calculated from the MaxQuant protein intensity results across the species or tissue subtype groups. Protein ranked abundances were calculated from the median abundances.

### Peptide-based model of protein quantification

The peptide-based model of proteomic analysis was performed using the R package MSqRob (robust statistical inference for quantitative LC-MS proteomics)^27,28^. Briefly, MSqRob adopts a linear mixed model framework and calculates protein intensity informed by all peptide intensities. In addition, MSqRob performs ridge regression, borrows information across proteins, and down-weighs outliers. MSqRob Shiny App v0.7.7 utilised the MaxQuant search output files and the parameters were followed as described in Goeminne *et al.* 2018^28^. Data underwent log_2_ transformation and quantile normalization as part of pre-processing. Explant subtype (non-implanted, non-degenerated, thrombus, neotissue, and calcification) were set as the fixed effect and donor, sample, and peptide sequence set as the random effects. Peptide intensities for dendrogram and heatmap analysis were extracted from the MSqRob data output and calculated as the overall median value of all the median peptide intensities for each individual peptide sequence, across either explant subtype or implantation duration, depending on the analysis.

### Cross-species network analysis

Median abundances were calculated from the MaxQuant protein intensity results across the tissue subtype groups. Protein vector matrices were then constructed with 25 distinct dimensions, comprising five potential species tags (bovine-specific, bovine ortholog, human ortholog, human-specific, and “cannot differentiate”) across five explanted BPV degenerated subtypes (non-implanted, non-degenerated, thrombus, neotissue, and calcified), to represent protein profiles across species and disease. Additional matrices were created with four dimensions (all species tags; non-implanted vs. non-degenerated, thrombus, neotissue, or calcified) to represent protein profiles considering species tags within a disease stage. For each protein, vector matrices were constructed across 25 or 4 dimensions, with binary values (0 = unidentified, 1 = identified) for each species-disease combination. The distance between each protein vector in our protein-protein interaction network was calculated and used to weigh the network. Hamming distance was used to determine protein dissimilarity independently across tissue segments and species, while Jaccard similarity coefficient assessed similarity across the 25 distinct dimensions. Dissimilarity and Similarity thresholds were set below or above 0.5 vector distance, respectively. Network visualizations were made using Gephi v0.10.1.

### Statistical Analyses

GraphPad Prism v10.1.2 was used for statistical analyses. Data are presented in boxplots, and two group comparisons were performed with Mann Whitney U tests. Heatmaps were generated from exported MSqRob peptide-level quantifications. ANOVA tests were performed in Perseus v2.1.3.0. One way ANOVA was used to test changes in peptide level intensity for bovine proteins. Two-way ANOVA was used to test changes in peptide level intensity for proteins across species, across tissue subtypes and the interaction between the two. To plot the heatmaps and perform dendrogram clustering, median peptide quant per tissue subtype sample were utilized. Rows were z-score normalized. All clustering was performed unbiasedly, except for implantation duration length of time was used to constrain the Euclidean distance calculations by increasing order of years. To compare the peptide intensities within tissue subtype between species 1) where there were no orthologous peptides quantified, no test was performed (NA), 2) where one orthologous peptide was quantified, all values were compared using either t-test or Mann Whitney U test, depending on the normality, and 3) where more than one orthologous peptide were quantified, the median peptide intensity was calculated for each peptide, and comparisons were performed using paired t-tests or Mann Whitney U tests, depending on the normality. Intra-species comparisons across segments were compared using repeated measures one-way ANOVA or Friedman’s test, depending on normality.

## Abbreviations

BPV: Bioprosthetic Valve
ECM: Extracellular Matrix
MS: Mass Spectrometry

## Acknowledgements

We acknowledge Boston Scientific Corporation for providing non-implanted glutaraldehyde-fixed bovine pericardial tissue for research use.

## Author contributions (CRediT)

Rachel Cahalane (Conceptualization, Data curation, Formal analysis, Funding acquisition, Writing – original draft)

Writing – review and editing)

Marie Billaud (Resources, Writing – review and editing)

Taku Kasai (Data curation, Writing – review and editing)

Amber Hendrickx (Data curation, Writing – review and editing)

Ludger J.E Goeminne (Writing – review and editing)

Jochen Muehlschlegel (Resources, Writing – review and editing)

Filip Rega (Resources, Writing – review and editing)

Masanori Aikawa (Resources, Writing – review and editing)

Laoise McNamara (Funding acquisition, Writing – review and editing)

Sasha Singh (Resources, Writing – review and editing)

Cassandra Clift (Conceptualization, Data curation, Formal analysis, Writing – original draft)

Elena Aikawa (Conceptualization, Funding acquisition, Writing – review and editing)

