## Supplemental Methods for "Cross-species proteomics quantification pipeline distinguishes donor versus host extracellular matrix in explanted biomaterials"

#### Supplementary Information


#### Authors

Rachel M.E. Cahalane[1,2], Bart Meuris[3], Joan T. Matamalas[1], Mark C. Blaser[1], Marie Billaud[4], Taku Kasai[1], Amber Hendrickx[3], Ludger J.E. Goeminne[5,6], Jochen Muehlschlegel[7], Filip Rega[3], Masanori Aikawa[1,4], Laoise McNamara[2,8], Sasha A. Singh[1,4], Cassandra L. Clift[1]* and Elena Aikawa[1,4]*

*Co-corresponding authors

#### Affiliations

[1]Center for Interdisciplinary Cardiovascular Sciences, Cardiovascular Division, Department of Medicine, Brigham and Women's Hospital, Harvard Medical School, Boston, MA, United States.

[2]Mechanobiology and Medical Device Research group (MMDRG), Biomedical Engineering, School of Engineering, College of Science and Engineering, University of Galway, Galway, Ireland.

[3]Department of Cardiovascular Sciences, KU Leuven, Leuven, Belgium.

[4]Center for Excellence in Vascular Biology, Cardiovascular Division, Department of Medicine, Brigham and Women's Hospital, Harvard Medical School, Boston, MA, United States.

[5]Gladyshev lab, Brigham and Women's Hospital, Harvard Medical School, Boston, MA, United States.

[6]Department of Anesthesiology and Critical Care Medicine, Johns Hopkins University School of Medicine, Baltimore, MD, United States.

[7]CÚRAM, SFI Research Centre for Medical Devices, Galway, Ireland.


**Short Title:** Cross-species bioprosthetic valve proteomics

### Supplemental Methods

#### Method Development; Collagenase Digestion and Antigen Retrieval

Approximately 1 x 1 cm control, non-implanted bioprosthetic tissue samples were processed into ~ $1mm^3$ pieces using a blade (n=8 in total). Four samples underwent collagenase digestion (one without and three with antigen retrieval steps, see below). The tissue pieces were incubated at 37°C for 1 hour in 1 mg/ml of filter-sterilized collagenase type IA-S from Clostridium histolyticum (Sigma-Aldrich C5894) in DMEM with gentle mixing. After 1 hour, the DMEM-collagenase solution is removed, and fresh DMEM-collagenase solution is added, with further gentle mixing at 37°C for 3 hours. Tissue pieces were then rinsed in PBS. Four samples underwent antigen retrieval steps only (one control and three antigen retrieval methods). Three antigen retrieval methods[44–46] were tested: 0.01M Citrate buffer (pH 6), 0.001M EDTA (pH 7.98), and 10mM Tris (pH 9.05). Tissue pieces were covered in the antigen retrieval solution or control (PBS). Antigen retrieval solutions and tissue pieces were heated to 95°C for 20 mins with no agitation. The PBS control was kept in the fridge for 20 mins. Buffer exchange was performed with 2 washes of 50mM ammonium bicarbonate. All tissue pieces were pulverized in liquid nitrogen using a pestle and mortar, and were then frozen at -20°C. Pulverised tissues were then resuspended in RIPA buffer (ThermoFisher) supplemented with protease and phosphatase inhibitors (Millipore Sigma), subjected to 10 mins sonication in a benchtop ultrasonic waterbath (Branson 5510), and followed by centrifugation at 2000 g for 5 min. The supernatant was collected and transferred to a new tube. The protein concentration was determined using the Bicinchoninic acid (BCA) assay on a Nanodrop spectrophotometer reading at 562 nm against a standard curve. All samples were prepared for mass spectrometry-based proteomics using 10µg of protein and the Preomics x96 iST kit according to the manufacturer’s guidelines, as described below. Ultimately loss of identifications with collagenase digestion and no gain in identifications with antigen retrieval steps, therefore neither collagenase digestion nor antigen retrieval steps were performed for the final sample processing (**Supplementary Fig. 9**)

#### Supplemental Figures

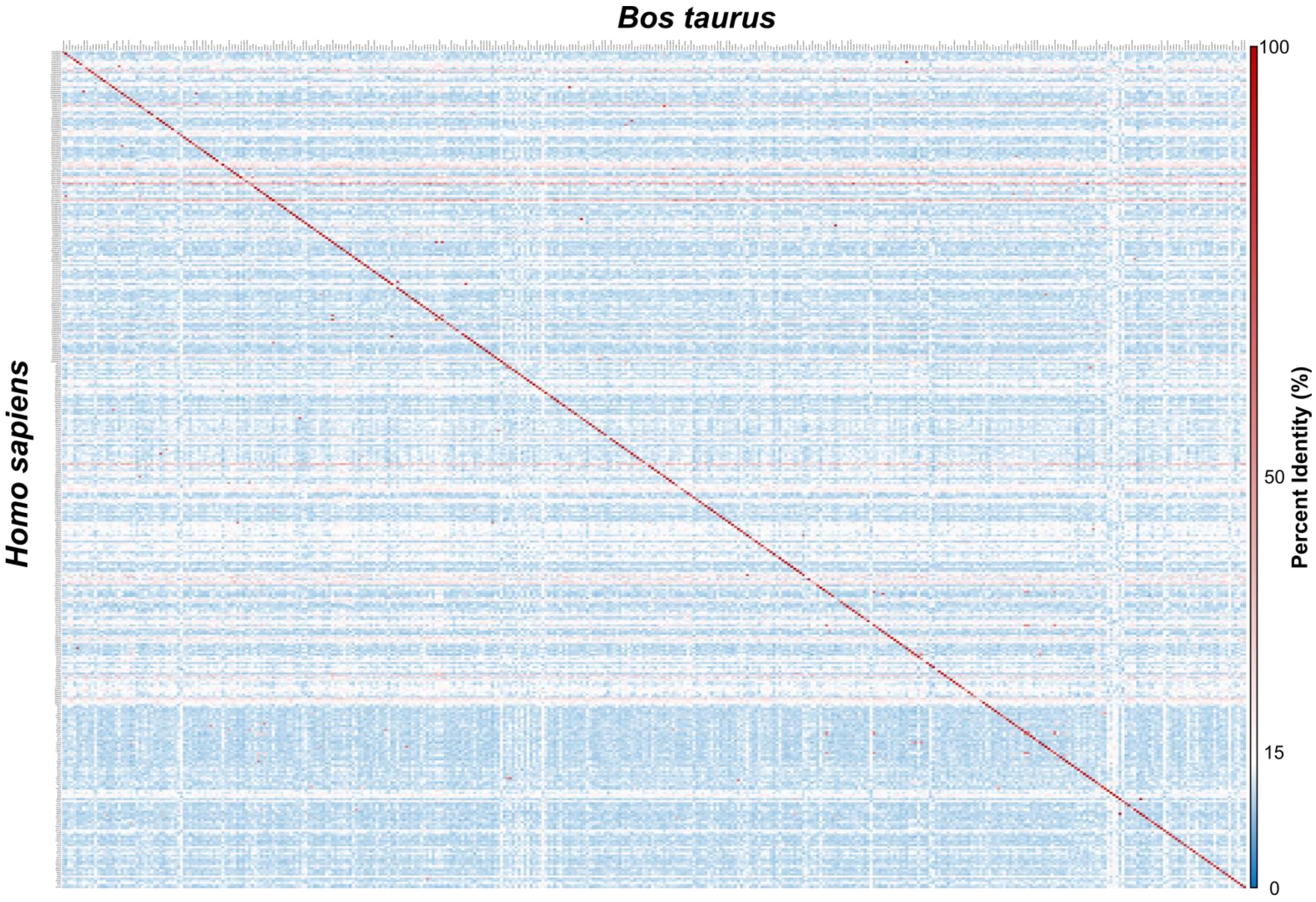


**Figure S1:** Percent identity matrix for custom *Bos Taurus* and full *Homo sapiens* n=400 potential overlapping proteins.

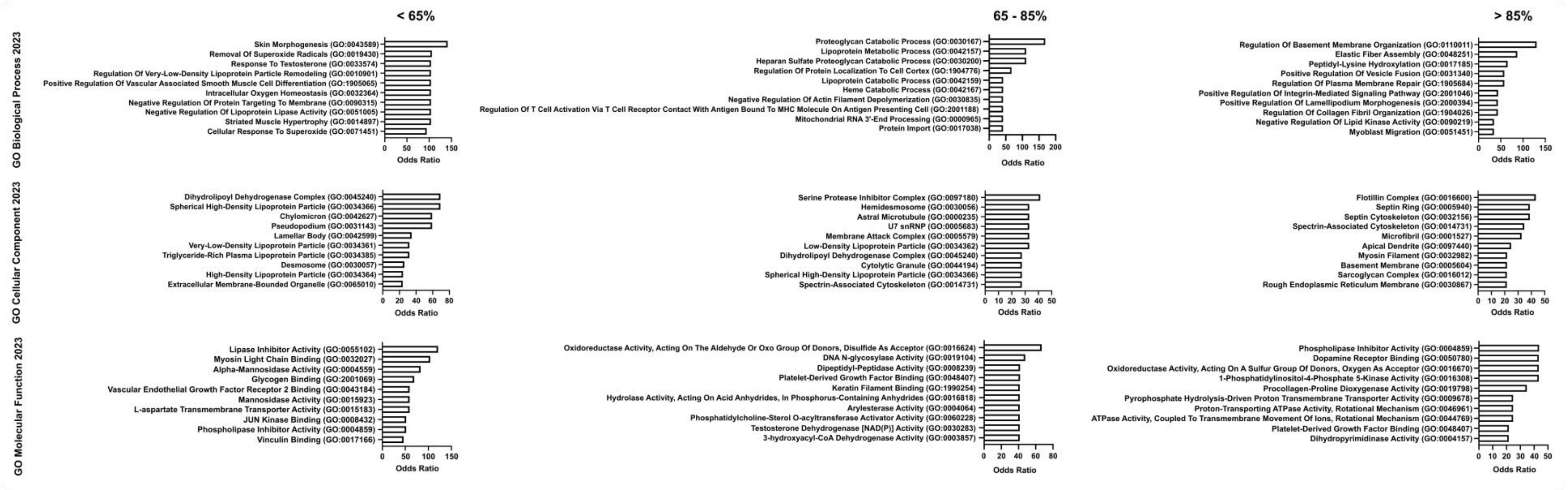

**Figure S2:** Top 10 Gene Ontology (GO) Biological Processes (top), Cellular Components (middle), and Molecular Function) for proteins with <65%, 65 - 85%, and >85% identity, sorted by odds ratio (from left to right, respectively)

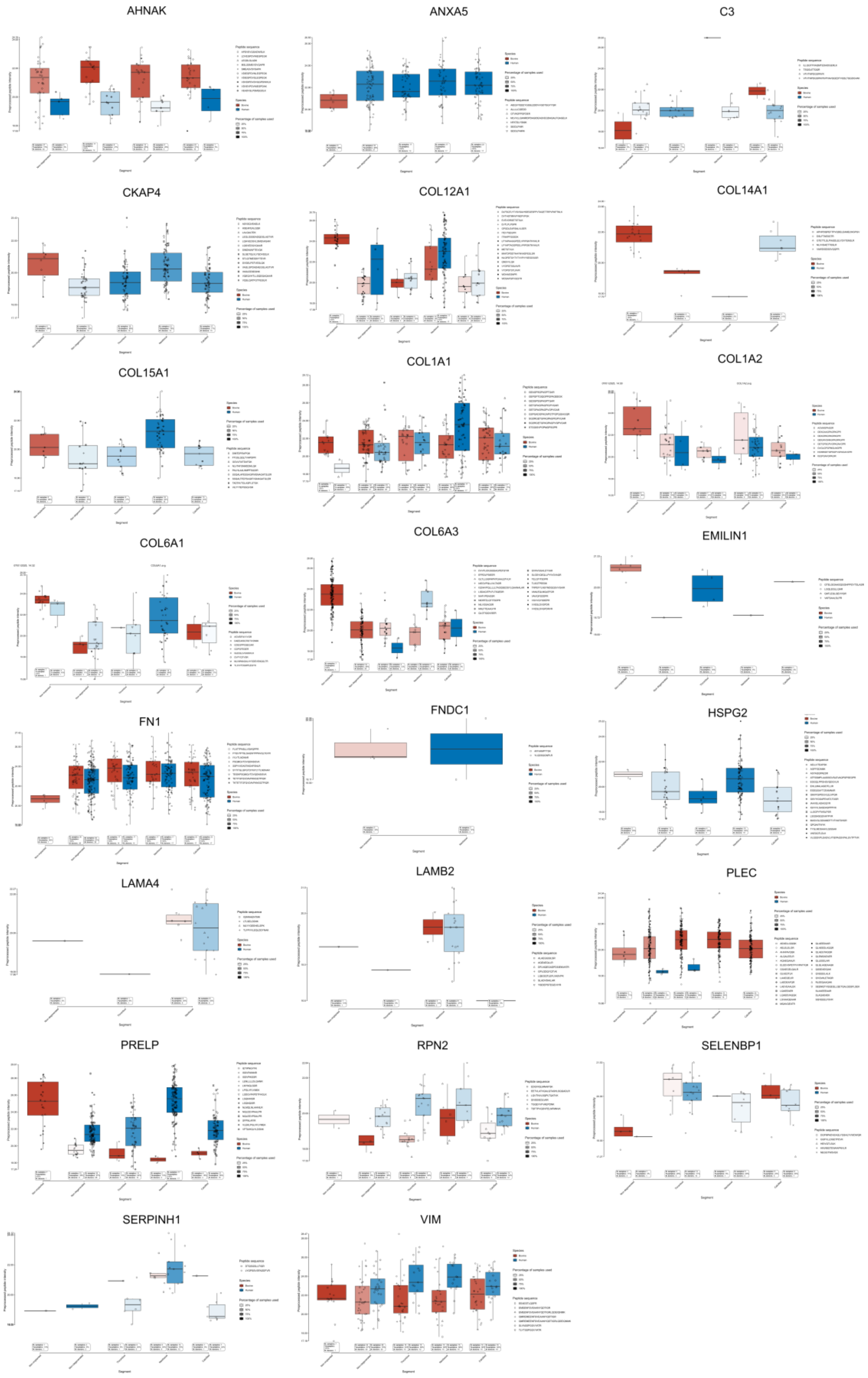


**Figure S3:** MSqRob peptide quantification boxplots of all orthologous proteins.

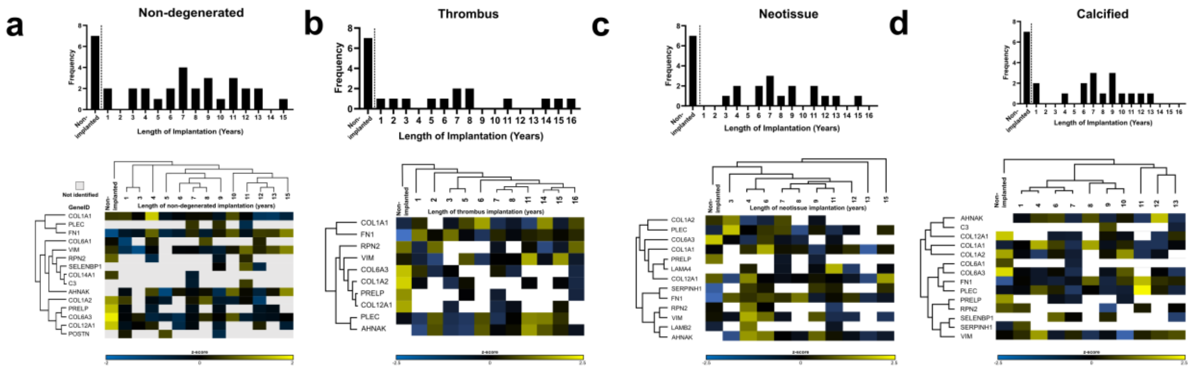


**Figure S4: Bovine protein quantification, plotted by length of BPV implantation.** Time per tissue subtype **a**, non-degenerated, **b**, thrombus, **c**, neotissue, and **d**, calcified. Hierarchical clusters are ordered on the x-axis, with Euclidean distance dendrograms shown.

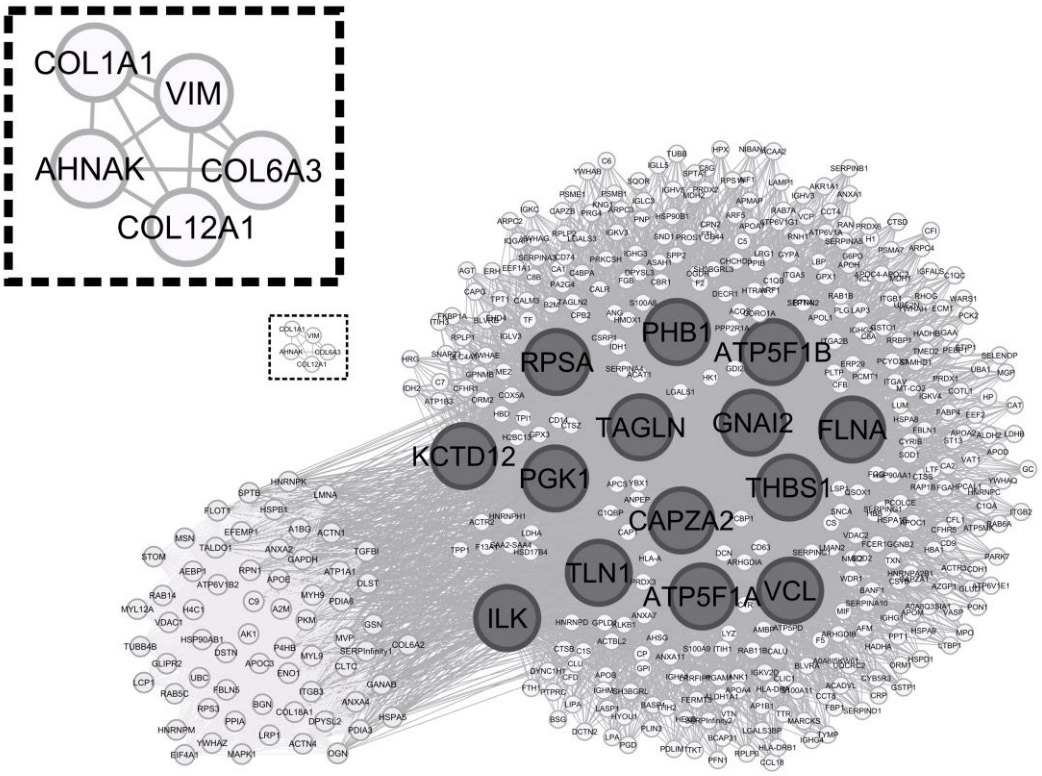


**Figure S5: Similarity-based metrics to prioritize proteins with largest dissimilarity.** Hamming distance vector analysis showing dissimilar proteins across degeneration subtype tags (non-implanted, non-degenerated, thrombus, neotissue, calcified), which were then used as edge weights in the protein similarity network. Dissimilarity thresholds were set below 0.5 vector distance.

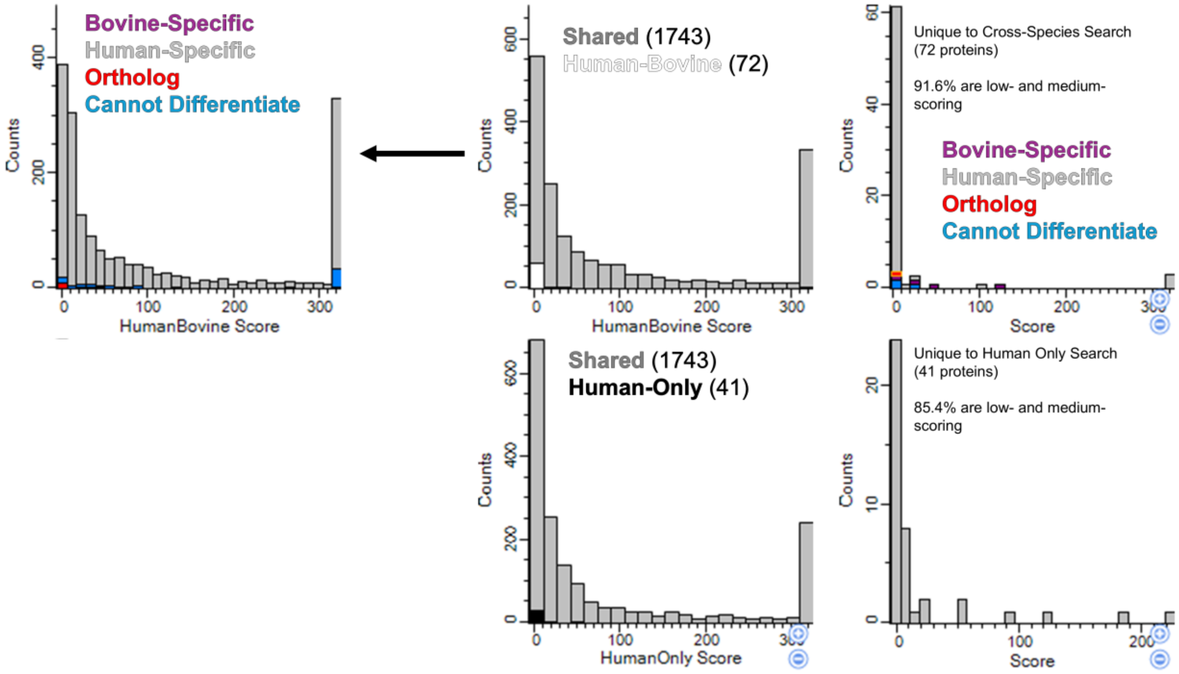


**Figure S6: Protein scoring.** Plotted protein confidence andromeda scores, comparing distribution to the cross species method (HumanBovine score) compared to the human FASTA only search strategy (HumanOnly score). Amongst all protein identifications, HumanOnly search strategy increases low to medium scoring by 10.62%. Unique proteins identified for each search are low to medium scoring: For HumanBovine: 91.6% of unique proteins are low/medium; HumanOnly: 85.4%. Andromeda score binning: Low score <10; medium 10-40; high >40.

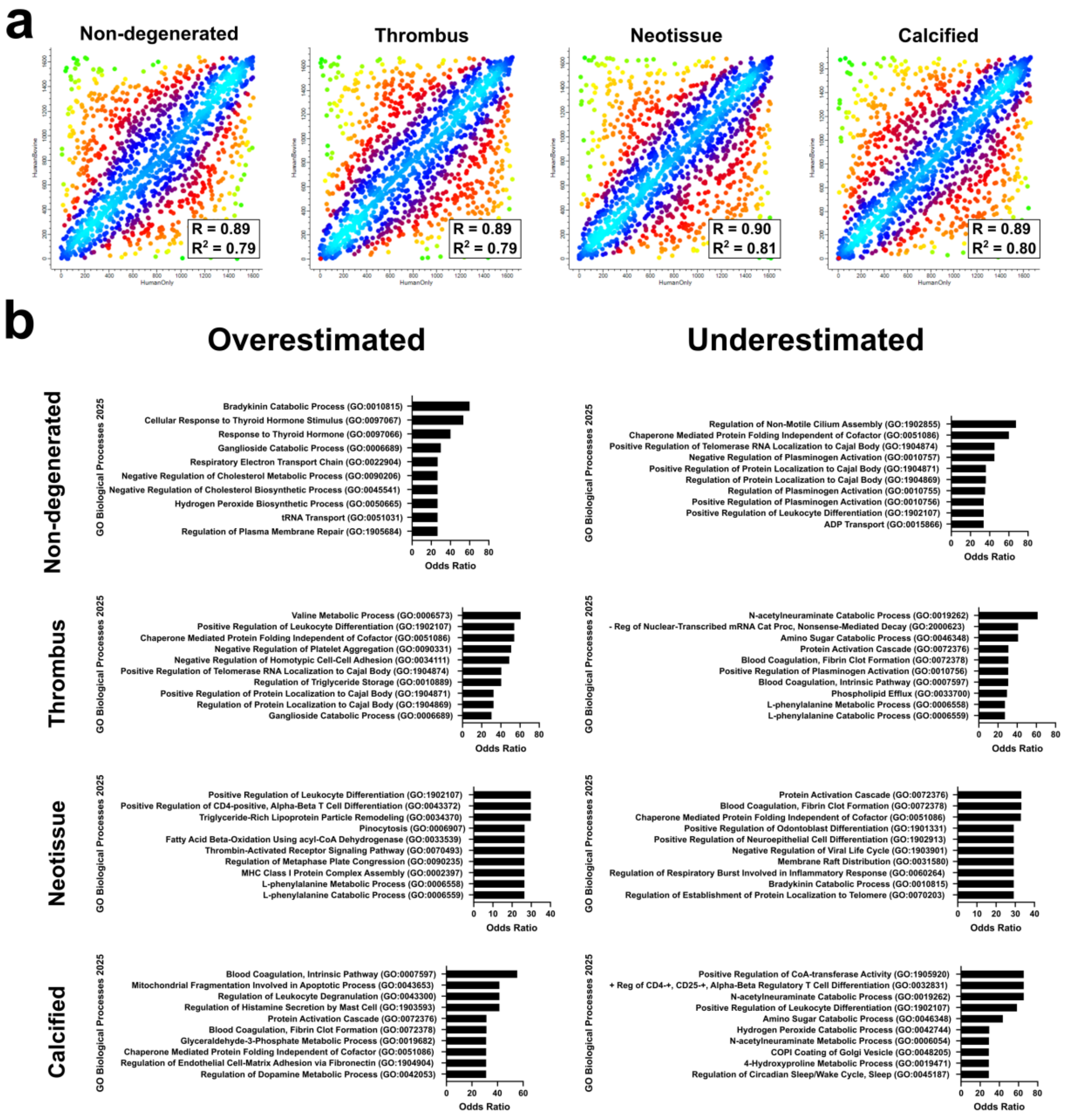


**Figure S7: Comparison of cross-species search results against a human only search.** **a.** Rank scatter density plots for non-degenerated, thrombus, neotissue, and calcified tissue subtypes. **b.** Top 10 Gene Ontology Biological Processes 2025 ranked by odds ratio for the over- and underestimated proteins per tissue subtype.

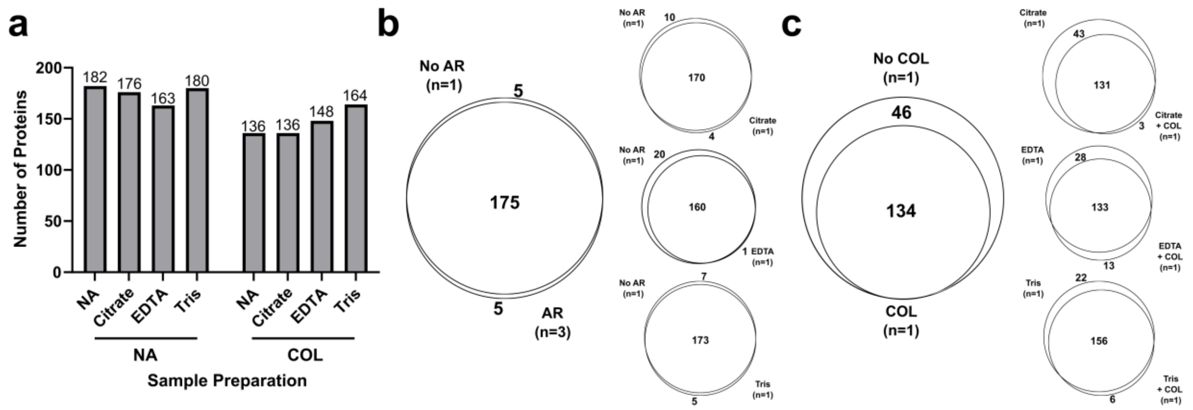


**Figure S8: Glutaraldehyde fixed tissue proteomics preparation method optimization. a**, Number of proteins identified with the different epitope retrieval preparation methods. **b**, Venn diagrams comparing antigen retrieval (AR) methods: none, citrate, EDTA and Tris. **c**, Venn diagrams comparing collagenase digestion (COL) methods, with or without subsequent antigen retrieval methods. NA = none.

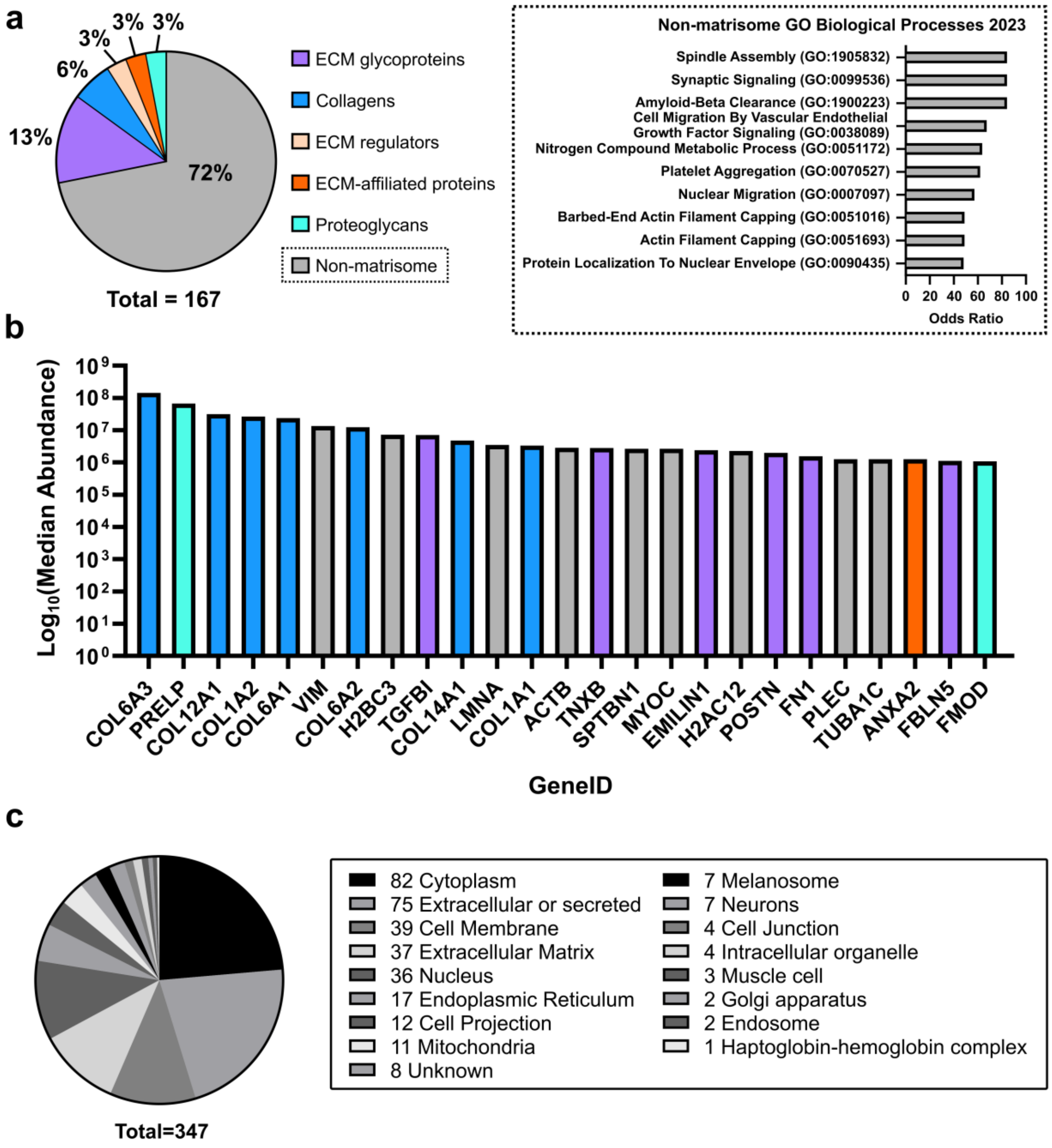


**Figure S9: Non-implanted control glutaraldehyde-fixed bovine pericardium proteome.** **a**, Identifications: Pie chart demonstrating the percent of identified proteins categorised according to the *Bos taurus* matrisome[1]. Non-matrisome (72%) GeneIDs summarized by the top 10 Gene Ontology (GO) Biological Processes 2023[2]. **b**, Quantifications: Top 25 proteins ranked by median total peptide normalized abundance (log scale) across the n=7 samples profiled (Proteome Discoverer v2.5). **c**, Pie charts demonstrating frequency of subcellular locations (n = 347) reported for the proteins (n = 167) using the UniProt database[3].

#### Supplemental Tables

**Table S1:** *Bos taurus* and *Homo sapiens* proteins with 100% identity.

| GeneID | Protein Description | Bovine Accession | Human Accession |
|---|---|---|---|
| SEPTIN7 | Septin-7 | F1MIH2 | Q16181, A0A024RA87, E7EPK1, E7ES33, G3V1Q4 |
| PIP4K2B | Phosphatidylinositol-5-phosphate 4-kinase type 2 beta | F1MHT2 | P78356 |
| FLOT1 | Flotillin-1 | Q08DN8 | O75955 |
| RPS18 | Small ribosomal subunit protein uS13 | Q3T0R1 | P62269, A0A0G2JQH2, Q5GGW2 |
| TNC | Tenascin C | A0A3Q1LLF1 | J3QSU6, E9PC84, F5H7V9 |
| RAB2A | Ras-related protein Rab-2A | A0A3Q1MJU0 | P61019, E9PKL7 |
| EEF1A2 | Elongation factor 1-alpha 2 | Q32PH8 | Q05639, A0A2U3TZH3, A0A9L9PXK0, A0A9L9PYI8, A0A6Q8PFK6 |
| RPL15 | Large ribosomal subunit protein eL15 | Q5EAD6 | P61313, E7ERA2 |
| RAB14 | Ras-related protein Rab-14 | Q3ZBG1 | P61106, A0A994J451, A0A994J4B9, A0A994J774 |
| YWHAZ | 14-3-3 protein zeta/delta | P63103 | P63104, B0AZS6, B7Z2E6 |
| UBC | Polyubiquitin-C | P0CH28 | P0CG48, Q96C32 |
| ARF3 | ADP-ribosylation factor 3 | Q5E9I6 | P61204 |
| SF3B3 | Splicing factor 3B subunit 3 | A0JN52 | Q15393 |
| PSMD11 | 26S proteasome non-ATPase regulatory subunit 11 | Q2KI42 | O00231 |
| PSA5 | Proteasome subunit alpha type-5 | Q5E987 | P28066 |
| LSM3 | U6 snRNA-associated Sm-like protein LSm3 | A0A452DIA6 | P62310 |
| RS28 | Small ribosomal subunit protein eS28 | Q56JX6 | P62857 |
| RS21 | Small ribosomal subunit protein eS21 | Q32PB8 | P63220 |
| TBB4B | Tubulin beta-4B chain | Q3MHM5 | P68371 |
| TBB2A | Tubulin beta chain | E1BJB1 | Q13885 |
| RMD5A | RING-type E3 ubiquitin transferase | E1BHI2 | Q9H871 |

**Table S2: Cannot Differentiate Gene Ontology Biological Processes 2023, only adjusted P-value significant presented.**

| Gene Ontology Biological Process 2023 Term | Genes | Adjusted P-value | Odds Ratio |
|---|---|---|---|
| Elastic Fiber Assembly (GO:0048251) | EFEMP2;FBLN5 | 0.013039911 | 118.5119048 |
| Positive Regulation Of Mitophagy (GO:1901526) | VDAC1;SLC25A5 | 0.013039911 | 118.5119048 |
| Establishment Of Golgi Localization (GO:0051683) | COPG1;YWHAZ | 0.015705695 | 94.8047619 |
| Epithelial Cell-Cell Adhesion (GO:0090136) | ITGB5;THBS4;VCL | 0.001525703 | 89.93674699 |
| Positive Regulation Of Establishment Of Protein Localization To Telomere (GO:1904851) | CCT3;TCP1 | 0.021246577 | 67.71088435 |
| Regulation Of Establishment Of Protein Localization To Telomere (GO:0070203) | CCT3;TCP1 | 0.021473557 | 59.24404762 |
| Regulation Of Protein Localization To Cajal Body (GO:1904869) | CCT3;TCP1 | 0.021473557 | 59.24404762 |
| Positive Regulation Of Lipid Localization (GO:1905954) | LRP1;LCP1 | 0.021473557 | 59.24404762 |
| Positive Regulation Of Protein Localization To Cajal Body (GO:1904871) | CCT3;TCP1 | 0.021473557 | 59.24404762 |
| Purine Ribonucleoside Triphosphate Biosynthetic Process (GO:0009206) | ATP5F1A;ATP5F1B | 0.021473557 | 59.24404762 |

**Table S3: Top 12 Human Only Gene Ontology Biological Processes 2023 sorted by odds ratio, and adjusted P-value significant only.**

| Term | Genes | Adjusted P-value | Odds Ratio |
|---|---|---|---|
| Positive Regulation Of Leukocyte Differentiation (GO:1902107) | LGALS3;XRCC6;LGALS1;PRKDC;NCKAP1L;HMGB1;LGALS9 | 2.06607E-06 | 128093 |
| Cyclooxygenase Pathway (GO:0019371) | CBR1;PTGIS;PTGES2;TBXAS1;PTGDS;PTGS2;PTGS1 | 1.0255E-05 | 75.61157025 |
| Negative Regulation Of Nitric Oxide Biosynthetic Process (GO:0045019) | PTGIS;ROCK2;CAV1;KHSRP;SIRPA;GLA | 7.50636E-05 | 64.77168142 |
| Negative Regulation Of Nitric Oxide Metabolic Process (GO:1904406) | PTGIS;ROCK2;CAV1;KHSRP;SIRPA;GLA | 7.50636E-05 | 64.77168142 |
| Neutrophil Degranulation (GO:0043312) | VAMP8;VAMP7;STXBP3;STXBP2 | 0.002913971 | 43.13022982 |
| Regulation Of Basement Membrane Organization (GO:0110011) | LAMA2;LAMB1;LAMC1;NID1 | 0.002913971 | 43.13022982 |
| Blood Coagulation, Fibrin Clot Formation (GO:0072378) | FGB;FGA;GP9;GP1BB;F12;FGG;F13A1;FBLN1;GP1BA;F13B;GP5 | 5.97449E-08 | 39.69546351 |
| Protein Activation Cascade (GO:0072376) | FGB;FGA;GP9;GP1BB;F12;FGG;F13A1;FBLN1;GP1BA;F13B;GP5 | 5.97449E-08 | 39.69546351 |
| Fibrinolysis (GO:0042730) | FGB;FGA;SERPINB2;FGG;SERPINE1;SERPINF2;GP1BA;PLG;F2 | 2.46911E-06 | 32.43971631 |
| Fructose 1,6-Bisphosphate Metabolic Process (GO:0030388) | PFKL;TIGAR;KHSRP;ALDOC;FBP1;PFKP | 0.000216768 | 32.3840708 |
| Negative Regulation Of Platelet Aggregation (GO:0090331) | CEACAM1;SERPINE2;PRKCD;CD9;ALOX12;PRKG1 | 0.000216768 | 32.3840708 |
| Formation Of Cytoplasmic Translation Initiation Complex (GO:0001732) | EIF3M;EIF3K;EIF3L;EIF3I;EIF3G;EIF3H;EIF3E;EIF3F;EIF3C;EIF3B | 1.03772E-06 | 27.04760497 |

**Table S4: MaxQuant contaminants file editing.** Twenty-five proteins were removed from the standard MaxQuant contaminants FASTA file to create the modified contaminant database used in the current study. Accession ID and description for the 25 removed proteins are presented.

| Accession ID | Species | Description |
|---|---|---|
| A2I7N0 | *Bos taurus* | SERPINA3-4 |
| A2I7N1 | *Bos taurus* | SERPINA3-5 |
| A2I7N3 | *Bos taurus* | SERPINA3-7 |
| ENSEMBL:ENSBTAP00000016046 | *Bos taurus* | Similar to fibulin-1 C isoform 1 |
| ENSEMBL:ENSBTAP00000032840 | *Bos taurus* | Similar to apolipoprotein B |
| P02672 | *Bos taurus* | Fibrinogen alpha chain precursor |
| P02676 | *Bos taurus* | Fibrinogen beta chain precursor |
| P04258 | *Bos taurus* | Similar to Collagen alpha 1(III) chain |
| P15497 | *Bos taurus* | Apolipoprotein A-I precursor |
| P31096 | *Bos taurus* | Osteopontin |
| P81644 | *Bos taurus* | Apolipoprotein A-II precursor |
| Q03247 | *Bos taurus* | Apolipoprotein E precursor |
| Q05443 | *Bos taurus* | Lumican precursor |
| Q1A7A4 | *Bos taurus* | Similar to complement component C5 |
| Q28194 | *Bos taurus* | Thrombospondin-1 |
| Q29RQ1 | *Bos taurus* | Complement component C7 precursor |
| Q2KIT0 | *Bos taurus* | Similar to collagen |
| Q2KJC7 | *Bos taurus* | Periostin |
| Q2UVX4 | *Bos taurus* | Complement C3 precursor |
| Q32PJ2 | *Bos taurus* | Apolipoprotein A-IV precursor |
| Q3KUS7 | *Bos taurus* | Complement factor B |
| Q3MHN2 | *Bos taurus* | Complement component C9 precursor |
| Q3Y5Z3 | *Bos taurus* | Adiponectin precursor |
| Q3ZBS7 | *Bos taurus* | Vitronectin |
| Q862S4 | *Bos taurus* | Similar to pro alpha 1(I) collagen (Fragment) |
